# Destabilizing evolutionary and eco-evolutionary feedbacks drive empirical eco-evolutionary cycles

**DOI:** 10.1101/488759

**Authors:** Michael H. Cortez, Swati Patel, Sebastian J. Schreiber

## Abstract

We develop a method to identify how ecological, evolutionary, and eco-evolutionary feedbacks influence system stability. We apply our method to nine empirically-parameterized eco-evolutionary models of exploiter-victim systems from the literature and identify which particular feedbacks cause some systems to converge to a steady state or to exhibit sustained oscillations. We find that ecological feedbacks involving the interactions between all species and evolutionary and eco-evolutionary feedbacks involving only the interactions between exploiter species (predators or pathogens) are typically stabilizing. In contrast, evolutionary and eco-evolutionary feedbacks involving the interactions between victim species (prey or hosts) are destabilizing more often than not. We also find that while eco-evolutionary feedbacks rarely altered system stability from what would be predicted from just ecological and evolutionary feedbacks, eco-evolutionary feedbacks have the potential to alter system stability at faster or slower speeds of evolution. As the number of empirical studies demonstrating eco-evolutionary feedbacks increases, we can continue to apply these methods to determine whether the patterns we observe are common in other empirical communities.

## Introduction

A fundamental problem in community ecology is understanding what factors influence system stability, e.g., whether a community converges to a steady state or exhibits cycles. Empirical and theoretical studies have shown that feedbacks between ecological and evolutionary processes, called eco-evolutionary feedbacks, can influence community stability and lead to different population-level dynamics (1; 2; 3; 4; 5; 6; 7). For example, experimental bacteria and virus-bacteria systems with demonstrated eco-evolutionary feedbacks converge to steady state (8; 9) whereas experimental rotifer-algae systems exhibit cycles (10; 11; 3; 12; 13).

Previous theoretical work has explored the (de)stabilizing effects ecological and evolutionary dynamics have on each other via eco-evolutionary feedbacks. In particular, ecological dynamics have the potential to stabilize unstable evolutionary dynamics or destabilize stable evolutionary dynamics (14; 2; 15). Similarly, evolutionary dynamics can stabilize or destabilize ecological dynamics (4; 5; 15). In general, stability of a whole system is influenced by the effects species’ densities have on the dynamics of population densities (ecological feedbacks), the effects species’ traits have on the dynamics of evolving traits (evolutionary feedbacks), and the effects population densities and evolving traits have on each other’s dynamics (eco-evolutionary feedbacks). Previous theoretical work (7; 15; 16; 17) has explored when these feedbacks have stabilizing versus destabilizing effects, and shown that the strengths of those effects increase or decrease with changes in the relative rates of ecological and evolutionary change. Specifically, stability of the whole system in the slow evolution limit is determined by ecological and eco-evolutionary feedbacks whereas stability of the whole system in the fast evolution limit is determined by evolutionary and eco-evolutionary feedbacks.

While these theoretical results identify many possible outcomes, it is not well understood which particular feedbacks are responsible for causing stable versus cyclic population dynamics in empirical systems. First, while the observed rates of ecological and evolutionary change are similar in the above empirical studies, most of the theory assumes ecological rates of change are either much faster or much slower than rates of evolutionary change. Second, because most systems are not identical in their composition of species and traits, it is unclear how to make comparisons across systems. Third, many empirical systems involve multiple interacting species and multiple evolving traits, but because much of the theory focuses on models with a small number of species and traits, it is difficult to apply the theory. Thus, we need new theoretical tools that can extend current theory and identify broadly the effects of ecological, evolutionary, and eco-evolutionary feedbacks while simultaneously pinpointing the importance of particular feedbacks.

Building on prior theoretical work (7; 15; 16), we develop a method using feedbacks defined in terms of the stability of a subsystem, i.e., the interactions and dynamics of a set of variables when all other variables are held fixed (e.g., the ecological subsystem defines the dynamics of all population densities when all population-level traits are held fixed). Our method identifies how the stabilities of complementary pairs of subsystems (e.g., ecological vs. evolutionary subsystems) at the equilibrium of the whole system and the interactions between them (e.g., the effects the evolutionary subsystem has on the ecological subsystem) influence the stability of the whole system. In addition to facilitating comparisons across systems, our method extends existing theory to systems with any number of species and evolving traits. We apply the method to nine models from the literature that are parameterized to empirical systems. We use the method to identify (i) the effects particular ecological, evolutionary, and eco-evolutionary feedbacks have on stability of the whole system, (ii) when eco-evolutionary feedbacks alter what one would predict about system stability from just ecological and evolutionary feedbacks, and (iii) how those effects are influenced by the relative speeds of ecology and evolution. Our results help explain why some systems exhibit periodic cycles while others converge to steady state.

## Methods

### Selecting parameterized eco-evolutionary models from published studies

To identify studies with parameterized eco-evolutionary models, we searched Web of Science and Google Scholar with keywords such as “eco-evolutionary dynamics” and “evolution & population dynamics”. Studies were selected only if they included models that were parameterized using empirical data and that described ecological and evolutionary dynamics. Here, ecological dynamics mean changes in population densities. Evolutionary dynamics mean either changes in a continuous trait (e.g., pathogen virulence) or the frequencies of different clonal types (e.g., defended and undefended clones). Three studies (18; 19; 20) were excluded because the models did not have coexistence equilibria with standing genetic variation in at least one population. In total, we identified 9 studies consisting of six predator-prey models, one intraguild predation model, and two host-pathogen models; see Table 1 for a summary. Multiple entries are listed in Table 1 for models with multiple parameterizations; Bolker et al. (21) is an exception because the results are identical for all four parameterizations. These nine studies represent all published studies known to the authors.

**Table 1:** Effects of complementary pairs of subsystems on system stability in parameterized models from the literature.

| Study | Evolving | System | Stabilities of complementary subsystem pairs* |  |  |  |  |
| --- | --- | --- | --- | --- | --- | --- | --- |
|  | Species | Behavior | Eco & Evo | Victim evo | Victim eco or eco-evo<sup>†</sup> | Exploiter Evo | Exploiter eco or eco-evo<sup>†</sup> |
| <u>Predator-prey</u> |  |  |  |  |  |  |  |
| Becks et al. (3) | Prey | Cyclic | S - U | U - S | U - S |  | S - U |
| Frickel et al. (9) | Both | Stable | S - S | S - S | S - S | S - S | S - S |
| Haafke et al. (13) | Both | Cyclic | U - U | U - U | U - S | S - U | S - U |
| Kasada et al. (26) | Prey | Stable | S - U | U - S | U - S |  | N - U |
| Wei et al. (34) Fig. 5a | Both | Stable | S - S | S - S | S - S | N - S | N - S |
| Fig. 5b | Both | Stable | S - S | S - U | U - S | N - S | N - U |
| Yoshida et al. (11; 10) | Prey | Cyclic | S - U | U - S | U - S |  | S - U |
| <u>Intraguild predation</u> |  |  |  |  |  |  |  |
| Hiltunen et al. (35)<sup>‡</sup> Fig. 2.1b | Basal prey | Cyclic | S - U | U - S | U - S |  | S - U |
| Fig. 2.1c | Basal prey | Cyclic | S - U | U - S | U - U |  | U - U |
| Fig. 2.1d | Basal prey | Cyclic | S - U | U - S | U - S |  | S - U |
| <u>Host-parasite</u> |  |  |  |  |  |  |  |
| Bolker et al. (21) | Parasite | Stable | S - S |  | S - S | S - S | S - S |
| Duffy et al. (27) | Host | Stable | S - S | S - S | S - S |  | U - S |

\* The first and second letters define how the subsystem listed in the column and its complementary subsystem, respectively, affect the stability of the whole system (S= stabilizing, U = destabilizing, N = neutral effect, Eco = ecological, Evo = evolutionary, Eco-evo = eco-evolutionary).
<sup>†</sup> In systems without victim (exploiter) evolution, the victim (exploiter) eco-evolutionary and ecological systems are the same.
<sup>‡</sup> Victim subsystems only involve the basal prey variables and exploiter subsystems involve the intraguild prey and intraguild predator variables.

### Method overview

Details about our method are given below and in appendices S1-S3. In short, we converted each model into a general form, computed the Jacobian, and evaluated it at the coexistence equilibrium point determined by the parameters in the original study. With the Jacobian, we determined the stabilities of the various subsystems, compared them to the stability of the whole system, and explored how our results depended on the speed of evolution.

### A general eco-evolutionary model

We converted all models into a general form that describes the changes in the densities of *n* species (*N*_1_, …, *N*_*n*_) and *m* population-level traits (*x*_1_, …, *x*_*m*_),

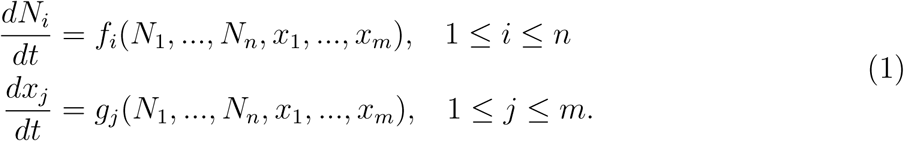

Here, *f*_*i*_ defines the ecological dynamics of species *i*; it accounts for all (possibly trait-dependent) intra and interspecific interactions involving species *i* (e.g., cooperation, competition, predation, and mutualism). The functions *g*_*j*_ define the evolutionary dynamics for each trait, which in general are density and frequency dependent. Note that clonal models with two clonal types (*C*_1_, *C*_2_) can be converted into continuous trait models by deriving equations for the total density (*N*_1_ = *C*_1_ +*C*_2_) and the frequency of clone 1 (*x*_1_ = *C*_1_*/N*_1_); see appendix S2 for additional details. Model (1) has been used previously to study equilibrium stability and species coexistence (15; 22). It encompasses other bodies of eco-evolutionary theory based on adaptive dynamics (23; 24) and quantitative genetics (25).

### Complimentary subsystem pairs and subsystem stability

We assume model (1) has a unique coexistence equilibrium where all species have positive densities; appendix S1 explains what changes when this assumption is not satisfied. We define stability of the whole system by the stability of the coexistence equilibrium, which is determined by the Jacobian (*J*), i.e., a derivative matrix that determines whether small perturbations from equilibrium decay (implying stability) or grow (implying instability). Mathematically, for stable systems all eigenvalues of the Jacobian have negative real parts and for unstable systems the Jacobian has at least one eigenvalue with positive real part. Importantly, each empirically parameterized model we considered has a unique coexistence equilibrium and if the coexistence equilibrium is unstable, then the system exhibits cycles because the equilibrium underwent a Hopf bifurcation.

Our method focuses on the stabilities of complementary pairs of subsystems. A subsystem describes the dynamics of a subset of variables when all other variables are fixed at their equilibrium values. Two subsystems form a complementary pair if together the subsystems include all variables in the system without overlap. For example, the (*n*-dimensional) ecological subsystem describes the population dynamics of all species (*dN*_*i*_*/dt* equations) when all traits are fixed at their equilibrium values (solid box in figure 1B). Its complement is the (*m*-dimensional) evolutionary subsystem (dashed box in figure 1B), which describes the evolutionary dynamics of all traits (*dx*_*j*_*/dt* equations) when all population densities are fixed at their equilibrium values. Alternatively, an eco-evolutionary subsystem (solid box in figure 1C) could be the population and trait dynamics associated with one species, say *N*_1_ and *x*_1_. Its complementary subsystem (dashed box in figure 1C) is the population and trait dynamics of the remaining species: *N*_2_, …, *N*_*n*_, *x*_2_, …, *x*_*m*_.

**Figure 1:**
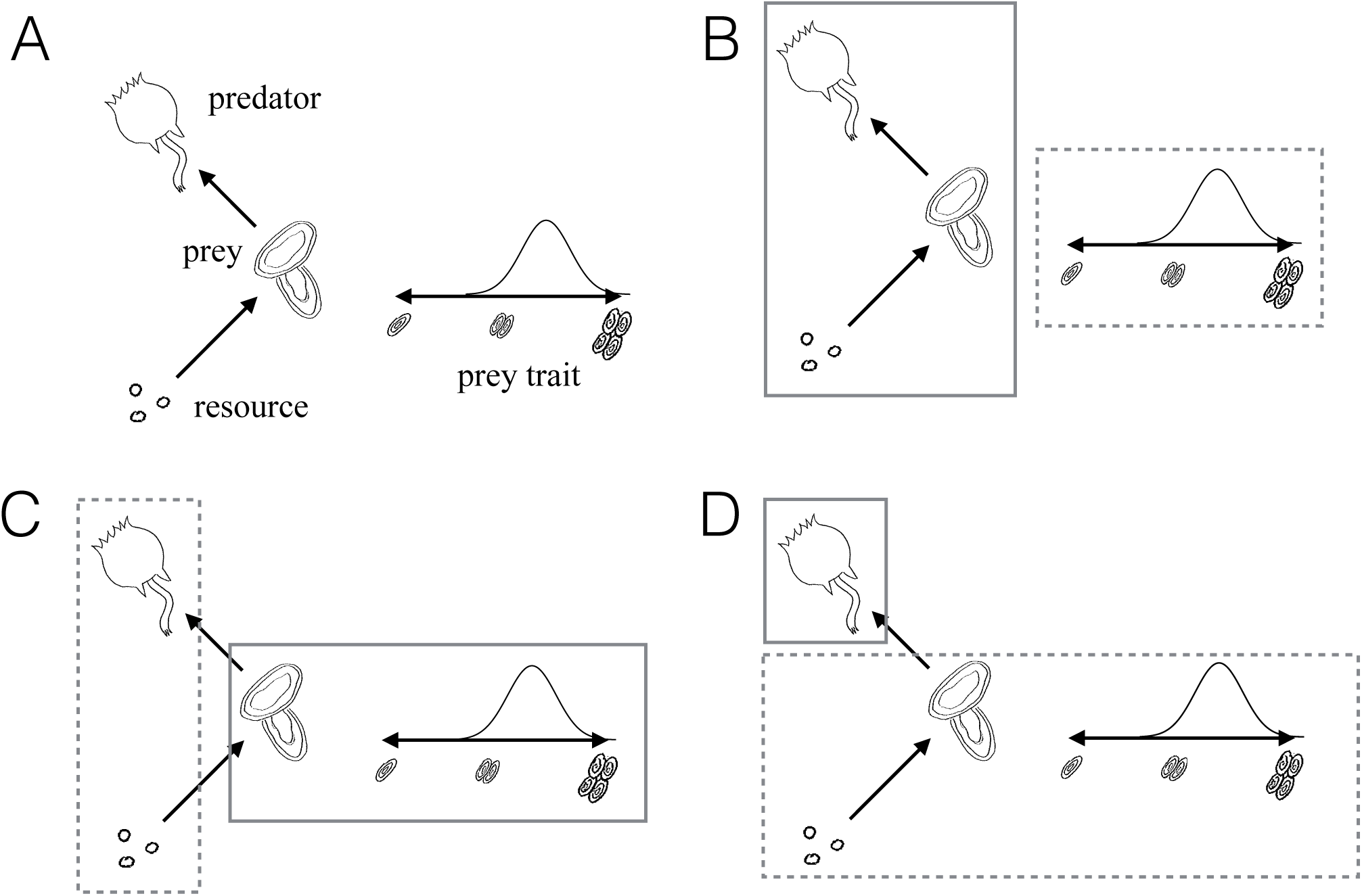
Examples of complementary subsystem pairs in the resource-prey-predator system with an evolving prey trait from Becks et al. (3). (A) The system dynamics involve changes in resource (nitrogen), prey (algae), and predator (rotifers) densities and the mean clump size of the prey. (B) Ecological subsystem (solid box) and its complementary evolutionary subsystem (dashed box). (C) Prey eco-evolutionary subsystem (solid box) and its complementary subsystem (dashed box). (D) Predator ecological subsystem (solid box) and its complementary subsystem (dashed box).

The stability of a subsystem is determined by the submatrix of the Jacobian that only involves the variables in that subsystem. For example, consider an eco-evolutionary nutrient-prey-predator model describing the dynamics of nutrient (*N*_1_), prey (*N*_2_), and predator (*N*_3_) densities and the mean prey trait (*x*_1_); this system is illustrated in figure 1. The Jacobian for this system has the form

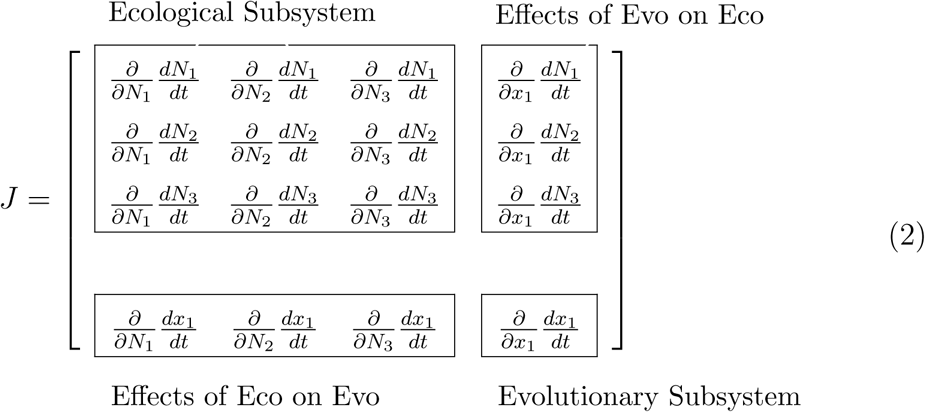

The top left box of the Jacobian determines the stability of the ecological subsystem (solid box in figure 1B), the bottom right box of the Jacobian determines the stability of the evolutionary subsystem (dashed box in figure 1B), and the off-diagonal boxes of the Jacobian determine the effects of ecology on evolution (bottom left) and the effects of evolution on ecology (top right). Mathematically, a subsystem is unstable if its submatrix has at least one eigenvalue with positive real part; a subsystem is stable if its submatrix has all eigenvalues with negative real parts; a subsystem is neutrally stable if its submatrix has all eigenvalues with non-positive real parts, at least one eigenvalue with strictly negative real part, and at least one eigenvalue with zero real part; and a subsystem is neutral if its submatrix has all eigenvalues with zero real parts; see figure S1 for illustrations of each type of stability.

### Stabilities of systems and their complimentary subsystem pairs

When there are no feedbacks between a pair of complementary subsystems, the stability of the whole system is determined by the stabilities of the complementary subsystems: the whole system is stable if both subsystems are stable and the whole system is unstable (implying cycles in our models) if either subsystem is unstable. When there are feedbacks between a pair of complementary subsystems, each subsystem has a stabilizing or destabilizing effect on the stability of whole system, but the feedbacks between the subsystems can alter the stability predicted by the complementary pair. For example, if the ecological subsystem is stable and the evolutionary subsystem is unstable in matrix (2), then the whole system is predicted to be unstable in the absence of eco-evolutionary feedbacks (zero entries in the top right or bottom left boxes). However, when eco-evolutionary feedbacks are present (non-zero entries in the top right and bottom left boxes) and stabilizing, the whole system can become stable. In this case, the feedbacks between the subsystems stabilize the whole system.

We consider four pairs of complementary systems, chosen for their biological relevance. First, the complementary ecological and evolutionary subsystems (figure 1B) identify the effects of ecological, evolutionary, and eco-evolutionary feedbacks involving all species. Second, the evolutionary subsystem for a single species (i.e, the subsystem composed of all evolving traits of one species) and its complement (also figure 1B) identify the effects of evolutionary feedbacks of a single species. Third, the eco-evolutionary subsystem for a single species (i.e., the subsystem composed of the density and all evolving traits for that species) and its complement (figure 1C) identify the effects of feedbacks within a single species. Fourth, the subsystem defined by all species and traits in a particular trophic level and its complement (figure 1D) identify the effects of feedbacks within a particular trophic level.

We use the stabilities of the complementary subsystem pairs to predict whether different feedbacks have stabilizing or destabilizing effects on the stability of the whole system in two ways. First, the stabilities of the complementary pairs of subsystems identify how subsystems affect the stability of the whole system. Specifically, unstable subsystems have destabilizing effects, stable or neutrally stable subsystems have stabilizing effects, and neutral subsystems have no direct effects on stability (but can indirectly affect stability through their interactions with other subsystems). See the appendix S1 for mathematical details and justifications.

Second, we compare the stabilities of the complementary subsystem pairs with the stability of the whole system in order to determine whether the feedbacks between subsystems do or do not alter system stability. There are four possibilities; the first and second correspond to cases where the feedbacks between complementary subsystems alter stability of the whole system from what would be predicted from just the stabilities of the complementary subsystems. First, if both subsystems are stable but the whole system is cyclic, then the feedbacks between the subsystems are destabilizing as they are sufficiently strong to counteract the stabilizing effects of the subsystems. Second, if one or both subsystems are unstable but the whole system is stable, then the feedbacks between the subsystems are stabilizing as they are sufficiently strong to counteract the destabilizing effects of the unstable subsystems. Third, if both subsystems are stable and the whole system is stable, then the feedbacks between the subsystems do not alter the stability of the system. Fourth, if one or both subsystems are unstable and the whole system is cyclic, then the feedbacks between the subsystems do not alter the stability of the system.

### Effects of varied evolutionary speed

To explore how the interactions between subsystem stability and the speed of evolution influence the stability of whole system, we varied the speed of evolution in the nine parameterized models. This was done by introducing multiplicative parameters into the right hand sides of the trait equations in model (1); see appendix S3 for details. We then assessed how speeding up and slowing down the rates of evolution influenced system stability and whether stable versus cyclic dynamics in the whole system could be accurately predicted from just the stabilities of the ecological and evolutionary subsystems.

## Results

### Effects of ecological, evolutionary and eco-evolutionary feedbacks on the stabilities of empirical systems

Across the nine parameterized models from the literature, subsystem stability differed depending on subsystem type (ecological, evolutionary, or eco-evolutionary) and species trophic level (exploiter vs. victim); see Table 1. Specifically, ecological subsystems were stable (or neutrally stable) in eight of the nine systems whereas evolutionary subsystems were stable in only four systems. Exploiter ecological, evolutionary, and eco-evolutionary subsystems were stable or neutral in seven systems. In contrast, victim ecological, evolutionary, and eco-evolutionary subsystems were stable in four systems.

With this information, we explored if feedbacks between subsystems altered the stability of the whole system from what would be predicted from just the stabilities of complementary subsystem pairs. What role do the feedbacks between subsystems play in influencing the stability of the four empirical systems that exhibit cycles (“cyclic” in column 4 of Table 1)? First, the evolutionary subsystem was unstable in all four systems and the complementary ecological subsystem was stable in three systems. This means that the feedbacks between the ecological and evolutionary subsystems were insufficiently strong to stabilize the system. Second, the evolutionary and eco-evolutionary subsystems for the victim species were unstable in all four systems and their complementary subsystems were stable in three systems. This means that the feedbacks between the victim subsystems and their complementary subsystems were insufficiently strong to stabilize the system. Third, the evolutionary, evolutionary, eco-evolutionary subsystems for the exploiter species were stable or neutral in three of the studies and their complementary subsystems were stable in all four systems. This means that the feedbacks between the exploiter subsystems and their complementary subsystems were destabilizing and sufficiently strong to alter the stability of the whole system.

What role do the feedbacks between subsystems play in influencing the stability of the five empirical systems that converge to equilibrium (“stable” in column 4 of Table 1)? First, in two systems, all subsystems we considered were stable (21; 9). This means that all feedbacks between the subsystems were either stabilizing or insufficiently strong to destabilize the whole system. Second, in three systems, there was at least one complementary subsystem pair made up of one stable and one unstable subsystem. For each of those systems, the feedbacks between the complementary subsystems were stabilizing and sufficiently strong to stabilize the whole system. For example, while the prey evolutionary and eco-evolutionary subsystems were unstable in Kasada et al. (26), the whole system was stable because the feedbacks between those subsystems and their complements were strongly stabilizing.

### Effects of evolutionary speed on stability

We explored how varying the speed of evolution affected system stability in the nine parameterized models. If varying the speed of evolution causes a change in stability, then it either causes a system undergoing cycles to converge to equilibrium or it causes a stable system to exhibit cycles; see appendix S3 for mathematical details. Varying the speed of evolution in the nine parameterized models produced one of four patterns (two shown in figure 2).

**Figure 2:**
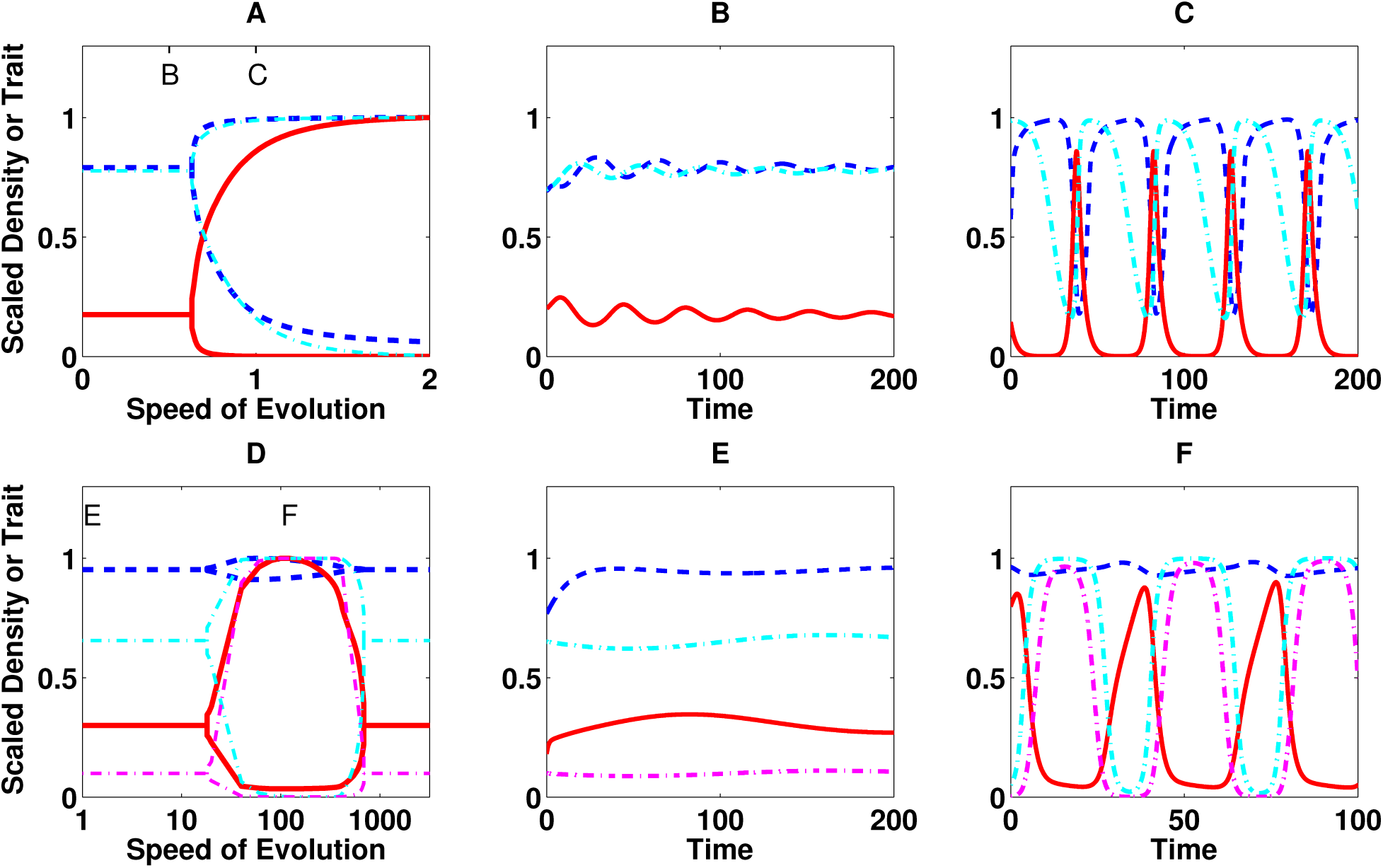
Predicted stability and dynamics of eco-evolutionary models with increased or decreased rates of evolution. (A-C) Dynamics of the Becks et al. (3) model with prey density (dashed blue), predator density (solid red), and proportion of defended prey (dash-dot cyan); nutrient dynamics are not shown. (D-F) Dynamics of the Duffy et al. (27) model with susceptible host density (dashed blue), infected host density (solid red), and proportions of resistant susceptible and infected hosts (dash-dot cyan and magenta, respectively). (A,D) Maximum and minimum long-term values for different evolutionary speeds; a single curve for each variable denotes the stable equilibrium value whereas two curves denote the maximum and minimum values during eco-evolutionary cycles. An evolutionary speed of one denotes the speed of evolution for the estimated parameter values in the original study. Letters denote evolutionary speeds for other panels.

First, for the four systems with stable ecological subsystems and unstable evolutionary subsystems (S-U in “Eco & Evo” column of Table 1), stability of the whole system switched from stable to unstable as the speed of evolution increased (figure 2A-C). In these systems, cyclic dynamics in the fast evolution limit are expected due to the instability of the evolutionary subsystem. Stability in the slow evolution limit is caused by stabilizing feedbacks between the ecological and evolutionary subsystems that are sufficiently strong to counteract the instability of the evolutionary subsystem. Hence, feedbacks between the ecological and evolutionary subsystems do not alter the stabilities of these systems in the fast evolution limit, but they do stabilize the systems in the slow evolution limit.

Second, for the Haafke et al. (13) study where the ecological and evolutionary subsystems were both unstable, the whole system exhibited cycles for all evolutionary speeds. The presence of cycles for all evolutionary speeds implies that the feedbacks between the ecological and evolutionary subsystems did not alter the stability of the system for any speed of evolution.

Third, for three of the four systems where the ecological and evolutionary subsystems were both stable (S-S in “Eco & Evo” column of Table 1), the whole system was stable for all evolutionary speeds. Stability for all evolutionary speeds implies that the feedbacks between the ecological and evolutionary subsystems did not alter the stability of any of the systems for any speed of evolution.

Fourth, the Duffy et al. (27) system, where the ecological and evolutionary subsystems were both stable, the whole system was stable for very fast and very slow evolutionary speeds and unstable for intermediate evolutionary speeds (figure 2D-F). Instability of the whole system for intermediate evolutionary rates means that the feedbacks between the ecological and evolutionary subsystems were sufficiently strong to destabilize the system only for intermediate speeds of evolution. A similar pattern has been observed in eco-evolutionary predator-prey models (7; 30; 16), but it is unclear if the same mechanisms are driving the pattern in the Duffy et al. (27) model because we lack general theory on when and why destabilization occurs at intermediate rates of evolution.

Overall, we found that the feedbacks between the ecological and evolutionary subsystems could alter the stability of the system at some evolutionary speed in five of the nine systems.

## Discussion

Our results identified that ecological, evolutionary, and eco-evolutionary feedbacks have systematically different effects on the stabilities of empirical systems and that those effects can depend on the species trophic level. Across the nine empirically parameterized models, ecological feedbacks tended to be stabilizing. In contrast, exploiter evolutionary feedbacks were stabilizing or neutral and victim evolutionary feedbacks were evenly split between stabilizing and destabilizing. Exploiter and victim ecological and eco-evolutionary feedbacks also consistently differed, with exploiter eco-evolutionary feedbacks being stabilizing or neutral and victim eco-evolutionary feedbacks being destabilizing more often than stabilizing. While our results are based on all empirically-parameterized models known to the authors, these models only represent a small number of systems, all of which involve exploiter-victim interactions. An important area of future work is applying and testing this theory in empirical systems with interactions other than exploiter-victim to understand whether ecological, evolutionary, and eco-evolutionary feedbacks have similar or different effects on stability in those systems. Our results help elucidate why some eco-evolutionary systems converge to steady state whereas others exhibit sustained cycles. (Recall that for our nine parameterized models, instability of the coexistence equilibrium implies cyclic dynamics.) The evolutionary subsystems were unstable in the four systems exhibiting cycles and stable in four of the five stable systems. This suggests that evolutionary feedbacks were important drivers of stability of our nine systems. In addition, in our models, instability and stability of evolutionary subsystems correspond to disruptive and stabilizing selection, respectively (2). Stabilizing and disruptive selection are observed with roughly equal frequencies across a broad set of empirical systems (28), suggesting that the destabilizing effects of evolutionary feedbacks are widespread across empirical systems.

Our results also help identify when eco-evolutionary feedbacks do and do not alter stability. First, in all but one system, the stability of the whole system could be predicted from just the stabilities of the ecological and evolutionary subsystems, implying eco-evolutionary feedbacks between all species did not alter the stability of the whole system. The one exception is the Kasada et al. (26) study, where we predict the eco-evolutionary feedbacks stabilized the whole system. Second, our results show that eco-evolutionary feedbacks involving just a subset of the species in the community could have different effects on stability. In particular, the eco-evolutionary feedbacks between the densities and traits of victim species could be stabilizing or destabilizing. This is consistent with prior theory predicting prey eco-evolutionary feedbacks can be stabilizing or destabilizing (2; 4; 7). In contrast, we found that the eco-evolutionary feedbacks between the densities and traits of exploiter species were stabilizing. Current theory predicts predator eco-evolutionary feedbacks can also be destabilizing (29; 30), but this was not observed in the four systems with exploiter evolution.

Our predictions about subsystem stability can be tested in empirical systems through controlled experiments in which some variables are held (nearly) fixed at their equilibrium values. One way to effectively fix evolutionary variables is to seed populations with lower standing genetic variation, e.g., as in (31; 3; 32; 33). If the magnitude of genetic variation is varied while the mean trait value is kept (effectively) constant, then the low genetic variation treatment will yield information about the stability of subsystems without that trait. Similarly, holding a species’ density nearly fixed will yield information about the stabilities of subsystems without that species. However, in most cases, subsystem stability cannot be determined by experiments where a variable is removed or changed substantially from its equilibrium value (e.g., removing a predator). This is because our subsystem-based approach assumes all fixed variables are held at their equilibrium values. It may be difficult or infeasible to hold densities or traits (nearly) constant in a given empirical system. Nonetheless, applying our theory to tailored, parameterized models allows one to make predictions about how specific feedbacks influence community stability and dynamics.

Our results highlight the need for additional theory to explain how the relative rates of evolution and ecology influence system stability. First, following Cortez (16), our approach can be extended to consider the effects of all subsystems. However, in systems with many species, the number of subsystems becomes very large, e.g, the Wei et al. (34) model with 10 variables has 1023 subsystems. Thus, new theory is needed to help understand what general rules govern how and when different subsystems influence system stability. Second, while current theory (7; 30; 16; 15) can explain model stability in the fast and slow evolution limits, we have a limited ability to make predictions about system stability when rates of ecology and evolution are similar. For example, it is unclear why the Duffy et al. (27) model exhibits cycles only at intermediate evolutionary speeds (figure 2D-F). This pattern has been observed in eco-evolutionary predator-prey models (7; 30; 16), but due to differences in subsystem stabilities and model dimension, it is unclear if the driving mechanisms are the same. Thus, theory is needed that explains how the speed of evolution interacts with subsystem stability to determine the stability of a whole system.

Our subsystem-based approach can be extended and potentially fruitful in other areas. First, applying our approach to a particular subsystem can help determine what feedbacks within that subsystem are responsible for its stability. For example, nearly all systems with unstable victim eco-evolutionary subsystems also had unstable victim evolutionary subsystems. Thus, instability of the eco-evolutionary subsystems must be due, in part, to the destabilizing effects of evolutionary feedbacks. Second, our approach may also be useful in purely ecological contexts. As examples, our approach could help identify how behavioral dynamics and species abundance dynamics affect community stability, how feedbacks within and between trophic levels affect the stability of food webs, how within-soil and above-soil communities contribute to the stability of plant-soil communities, and how environmental dynamics and species abundance dynamics affect system stability.

## Acknowledgements

The authors thank S.P. Ellner for sharing code. SJS was supported in part by the U.S. National Science Foundation grants DMS-1313418 and DMS-1716803.

## Data Accessibility

Appendices uploaded as online supplementary materials.

Computational files: zenodo DOI 10.5281/zenodo.3530691

## Supplementary Information

### S1 Notation, subsystems, and stability

Let 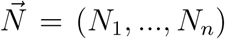 be the vector of species densities and 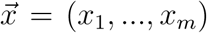 be the vector of evolving traits. Let 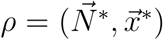 denote a coexistence equilibrium of the model from the main text, i.e., an equilibrium where all species have nonzero densities. When evaluated at *ρ*, the Jacobian of the model has the form

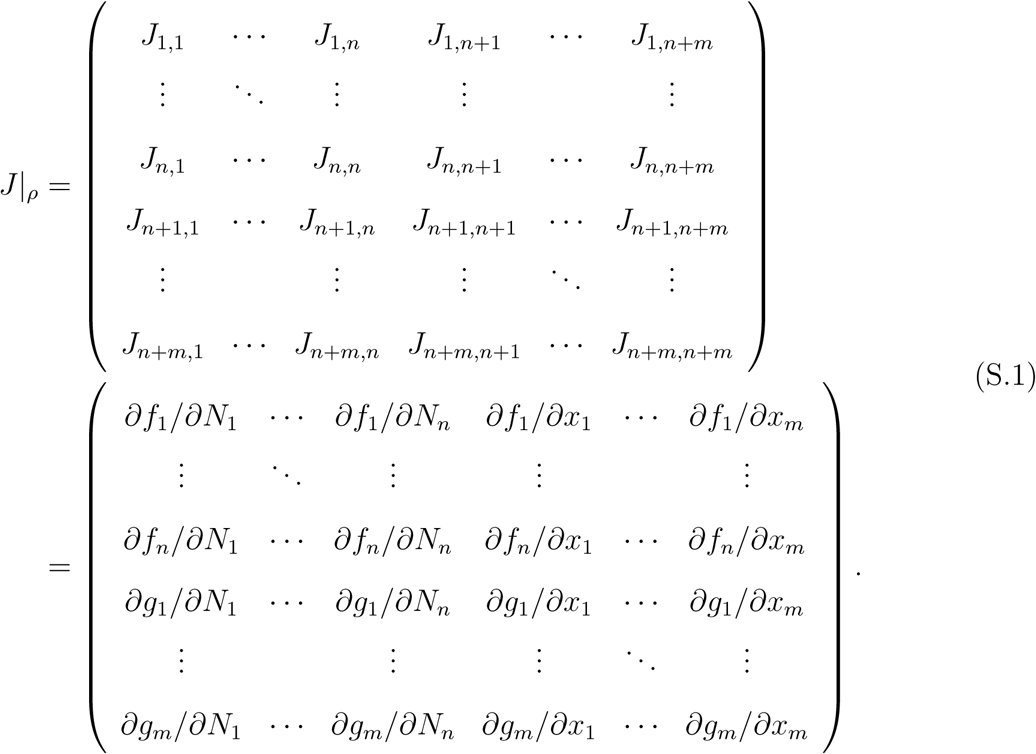

The equilibrium is stable if all eigenvalues of *J* have negative real parts, neutrally stable if all eigenvalues have non-positive real parts and at least one eigenvalue has negative real part, neutral if all eigenvalues have zero real parts, and unstable if at least one eigenvalue has positive real part. See figure S1 for an illustration of these definitions.

Let *S* be a subsystem containing variables {*s*_1_, *s*_2_, *…, s*_*k*_}. Let *M*_*S*_ be the submatrix of *J* made up of all entries 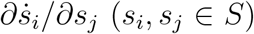. The stability of subsystem *S* when all variables not contained in *S* are fixed at their equilibrium values (defined by *ρ*) is determined by the eigenvalues of *M*_*S*_ following the same rules used for the coexistence equilibrium. Ecological subsystems contain only ecological variables (*N*_*i*_). Evolutionary subsystems contain only evolutionary variables (*x*_*j*_). Eco-evolutionary variables contain ecological and evolutionary variables. Two subsystems *S*_1_ and *S*_2_ form a complementary pair if their intersection is the empty set and their union contains all variables in the system.

**Figure S1:**
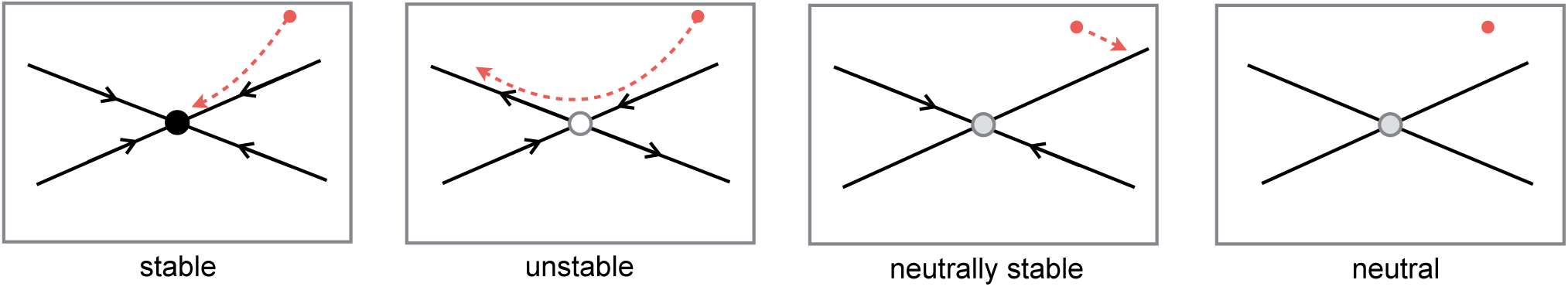
Two-dimensional examples of stable, unstable, neutrally stable, and neutral equilibria. Black lines are the eigenvectors computed from the Jacobian evaluated at the equilibrium. Black arrows denote the direction of the flow along the eigenvectors; an eigenvector with no arrow means no flow in that direction. Red dashed curves are example trajectories starting at the red dots.

In general, the stabilities of the subsystems influence the stability of the coexistence equilibrium. We begin by illustrating this in a two-dimensional system and then show how the idea extends to higher-dimensions. Consider a model with one ecological variable (*N*) and one evolutionary variable (*x*). Such a model has a 2×2 Jacobian (*J*) where the stability of the ecological subsystem is determined by the *J*_11_ entry, the stability of the evolutionary subsystem is determined by the *J*_22_ entry, and the interactions between the subsystems are defined by the product *J*_12_*J*_21_. When *J*_11_ < 0, the ecological subsystem has a stabilizing effect on the stability of the equilibrium and when *J*_11_ > 0, the ecological subsystem has a destabilizing effect on the stability of the equilibrium. Similarly, when *J*_22_ < 0 or *J*_22_ > 0, the evolutionary subsystem has a stabilizing or destabilizing effect, respectively, on the stability of the equilibrium. In the absence of bidirectional feedbacks between the ecological and evolutionary subsystems (i.e., *J*_12_*J*_21_ = 0), the stability of the equilibrium is determined by the stability of the ecological and evolutionary subsystems. However, when bidirectional feedbacks are present, the interactions between the two subsystems have stabilizing effects when *J*_12_*J*_21_ > 0 and destabilizing effects when *J*_12_*J*_21_ < 0.

A similar idea holds in higher dimensional models, where the entries *J*_11_ and *J*_22_ are replaced by submatrices of the Jacobian corresponding to a pair of complementary subsystems. The entries *J*_12_ and *J*_21_ are replaced by submatrices that represent the effects variables in one subsystem have on the dynamics of the variables in the other subsystem. Thus, stability of an equilibrium is affected by the stability of each subsystem as well as the feedbacks between the subsystems.

To show how stable, neutrally stable, neutral, and unstable subsystems influence equilibrium stability in higher dimensional models, we use use the characteristic polynomial of the Jacobian. As an example consider a four dimensional system with variables *N*_1_, *N*_2_, *N*_3_ and *x*_1_. The characteristic polynomial of the Jacobian (*J*) for such a system is

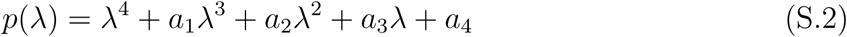

where

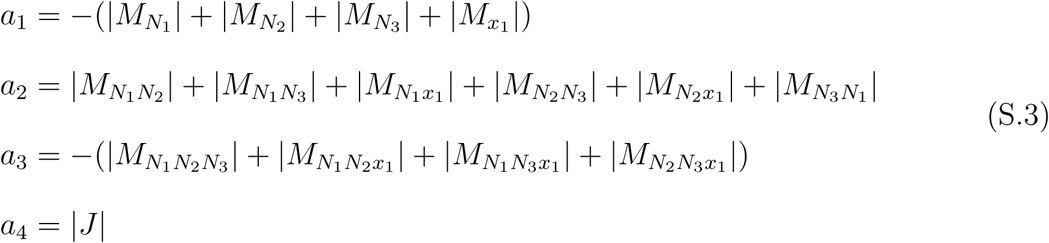

and | · | denotes the determinant of a (sub)matrix. A necessary condition for equilibrium stability is all coefficients of the characteristic polynomial are positive, i.e., *a*_*i*_ > 0 for all *i*. To see how the ecological subsystem (i.e., the *N*_1_, *N*_2_, *N*_3_-subsystem) affects the stability of an equilibrium, consider the characteristic polynomial for the submatrix 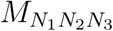,

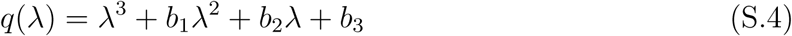

where

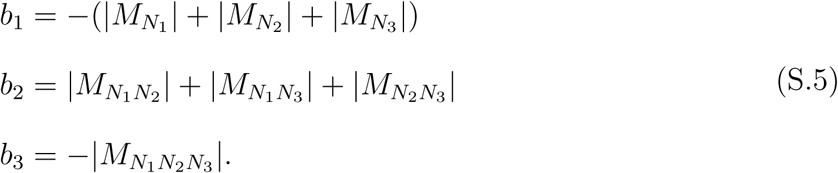

A necessary condition for stability of the ecological subsystem is *b*_*i*_ > 0 for all *i*. Notice that each term in the *b*_1_ equation also shows up in the equation for *a*_1_. More generally, *b*_*i*_ is a partial sum of the terms that define *a*_*i*_.

If the ecological subsystem is unstable, then at least one of the *b*_*i*_ coefficients is negative. Because *b*_*i*_ is a partial sum of the terms that define *a*_*i*_, this means *a*_*i*_ will be a more negative (or less positive) value. In terms of satisfying the necessary condition for equilibrium stability (*a*_*i*_ > 0), this means that an unstable ecological subsystem has a destabilizing effect on the stability of the equilibrium. In contrast, stable ecological subsystems (*b*_*i*_ > 0 for all *i*), neutrally stable ecological subsystems (*b*_*i*_ ≥ 0 for all *i*, with at least one of the *b*_*i*_ coefficients being positive), or neutral ecological subsystems with purely imaginary eigenvalues (also *b*_*i*_ ≥ 0 for all *i*, with at least one of the *b*_*i*_ coefficients being positive) have stabilizing effects on the whole system because the *a*_*i*_ coefficients are more positive if some of the *b*_*i*_ are positive. For neutral ecological subsystems where all eigenvalues are zero (*b*_*i*_ = 0 for all *i*), the subsystem has no direct effect on the stability of the whole system. For all of the models we examined, every neutral subsystem only had zero eigenvalues.

The above argument holds for a Jacobian of any size and a submatrix of any size. It explains our interpretation of how subsystems with different stabilities affect equilibrium stability. The above only focuses on the necessary condition for stability, i.e., all coefficients of the characteristic polynomial are positive. Necessary and sufficient conditions for stability are defined by the Routh-Hurwitz criteria. We do not use all of the conditions from the Routh-Hurwitz criteria because the other conditions are more complex and less tractable for large matrices.

### S2 Complementary subsystem pairs method

Here we detail how one can apply our complementary subsystem method to their own mode. Note that Step 1 is skipped for models with continuous traits.

#### Step 1: Convert discrete trait model into continuous trait model

For clonal models without stage structure where *C*_*i*_ is the density of clone *i*, the ecological variable of the continuous trait model is total density, *N* = ∑_*i*_ *C*_*i*_, and the evolutionary variables are the frequencies of clone *i, x*_*i*_ = *C*_*i*_*/N*, for *i* < *k*. For clonal models with stage structure where 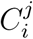 is the density of clone *i* in stage *j*, the ecological variables are the total densities in each stage, 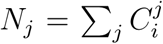, and the evolutionary variables are the proportions of clone *i* in each stage, 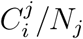 for *i* > 1. The differential equations for the continuous trait model are derived using the chain rule, e.g., 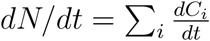 and 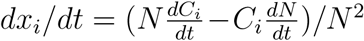 define the dynamics for a continuous trait model derived from a clonal model without stage structure.

#### Step 2: Find coexistence equilibria and determine their stability

For each coexistence equilibrium, *ρ*, stability is determined by computing the eigenvalues of the Jacobian evaluated at *ρ*.

#### Step 3: Pick a complementary subsystem pair and determine stabilities of the subsystems

Partition the state variables into two complementary. Let *S*_1_ be the subsystem containing variables from the first subset and *S*_2_ be the subsystem containing variables from the second subset. Find the two submatrices of the Jacobian corresponding to the two subsystems. In particular, the submatrix corresponding to subsystem *S*_1_ is made up of all entries 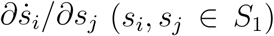 and the submatrix corresponding to subsystem *S*_2_ is made up of all entries 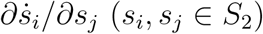. For each coexistence equilibrium, evaluate the two submatrices at the equilibrium and determine the stability of each subsystem by finding the eigenvalues of each submatrix.

#### Step 4: Compare stabilities of whole system and subsystem pairs

For each coexistence equilibrium, compare its stability with the stability of the complementary subsystems. If (i) both subsystems are stable and the coexistence equilibrium is unstable, (ii) both subsystems are unstable and the coexistence equilibrium is stable, or (iii) one subsystem is unstable, one subsystem is stable, and the coexistence equilibrium is stable, then the stability of the coexistence equilibrium differs from what is predicted from just the stabilities of the complementary subsystems. In such cases, the interactions between the subsystems alter the stability of the coexistence equilibrium based on what is predicted from just the stabilities of the complementary subsystems. In all other cases, the interactions between subsystems do not alter the stability of the coexistence equilibrium based on what is predicted from just the stabilities of the complementary subsystems.

We note two things about applying our method. First, applying our method to an unstable equilibrium (i.e., an equilibrium whose Jacocbian has at least one eigenvalue with positive real part) will identify which feedbacks are destabilizing that particular equilibrium. However, it may not explain why cyclic dynamics occur in that system, e.g., when there are unstable equilibria with homoclinic or heteroclinic orbits or cycles that arose via bifurcations not involving equilibria. Information about equilibrium instability is informative about cyclic dynamics only if the equilibrium has undergone a Hopf bifurcation that gave rise to an attracting periodic orbit (as was the case for the empirically parameterized models we considered).

Second, applying our method to a coexistence equilibrium always identifies how feedbacks affect stability at that specific equilibrium. However, if multiple equilibria are present, the feedbacks may have different effects at different equilibria. This is expected because the trait values and densities differ between the equilibria. However, it means that judgment must be used to determine which equilibria will yield biologically informative information. Applying our method to a stable coexistence equilibrium is always biologically informative because it identifies which feedbacks are responsible for stabilizing the system near that equilibrium. For unstable equilibria, our results are biologically informative if (i) the unstable equilibrium and an attracting periodic orbit are both present for the given parameter values and (ii) the unstable manifold of the equilibrium point intersects the stable manifold of the periodic orbit. In this case, our results help explain which feedbacks are responsible for destabilizing the unstable equilibrium and causing the system to exhibit periodic cycles. In other cases, e.g., when there are no periodic orbits or the equilibria have homoclinic or heteroclinic connections, applying our method to unstable equilibria may not explain system behavior.

### S3 Varying evolutionary speeds

To vary the speed of evolution in the models, we multiplied the right hand sides of the trait equations by the parameter *ϵ*_*i*_, where the rate of evolution is unchanged for *ϵ*_*i*_ = 1 and is slowed down (sped up) for 0 < *ϵ*_*i*_ < 1 (*ϵ*_*i*_ > 1). For example, the trait dynamics *dx*_1_*/dt* = *g*(·) are changed to *dx*_1_*/dt* = *ϵ*_1_*g*(·). For systems where a single species had multiple traits, each trait equation was multiplied by the same parameter (e.g., *ϵ*_1_ for all trait equations). For systems with multiple evolving species, the trait equations for different species were multiplied by different parameters (e.g., *ϵ*_1_ for the prey trait and *ϵ*_2_ for the predator trait). Our analysis only focused on the effects of varying each *ϵ*_*i*_ parameter independently; we did not explore how simultaneously varying multiple *ϵ*_*i*_ parameters would affect system stability.

Multiplying a trait equation by *ϵ*_*i*_ results in each entry of the Jacobian corresponding to that trait equation being multiplied by *ϵ*_*i*_. Thus, varying *ϵ*_*i*_ changes the magnitudes of those Jacobian entries, which in turn affects the eigenvalues of the Jacobian. For all of the models we considered, changing the rate of evolution (i.e., varying *ϵ*_*i*_) does not change the location of an equilibrium point. Thus, varying *ϵ*_*i*_ does not affect the entries of the Jacobian corresponding to the ecological equations or other trait equations.

For all models we considered, equilibria can only undergo Hopf bifurcations as the *ϵ*_*I*_ parameters are varied (15), and they cannot undergo bifurcations where one or more eigen-values are identically zero. The mathematical justification is the following. The determinant of the Jacobian factors as 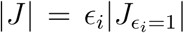 where 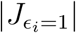 is the determinant of the Jacobian when *ϵ*_*i*_ = 1. Recall that the determinant of a matrix is equal to the product of its eigenvalues. If the equilibrium were to undergo a bifurcation with a zero eigenvalue, then 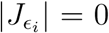 for some *ϵ*_*i*_ value, which would imply 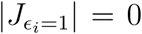 and further that 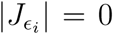 for all *ϵ*_*i*_ > 0. Thus, the sign of |*J* | is constant for *ϵ*_*i*_ > 0. This means that the eigenvalues of |*J* | can only change signs via Hopf bifurcations. In total, increasing or decreasing the rate of evolution can cause no effect on stability or cause the periodic orbit to undergo a Hopf bifurcation, i.e., either cause a system undergoing oscillations to converge to an equilibrium or cause a stable system to exhibit cycles.

### S4 Analysis of parameterized models

Here, we analyze the nine empirically parameterized models from the literature; subsystem stabilities are given in table S1. All calculations can be reproduced using the accompanying Maple worksheet and R files. However, the results for subsystem stability differ between the Maple worksheet and R files because of small numerical errors in R that result in R computing non-zero values for Jacobian entries that are analytically zero. If these numerical errors are accounted for (i.e., the entries are set equal to zero), then the results from R and Maple agree. In the following, Jacobian entries are listed as zero only if it is possible to show analytically that the entry is zero (see Maple worksheet for details). All other entries are rounded to three significant digits.

**Table S1:** Stabilities of complementary pairs of subsystems in parameterized models from the literature.

| Study | Evolving | System | Stabilities of complementary subsystem pairs* |  |  |  |  |
| --- | --- | --- | --- | --- | --- | --- | --- |
|  | Species | Behavior | Eco & Evo | Prey evo | Prey eco or eco-evo<sup>†</sup> | Pred Evo | Pred eco or eco-evo<sup>†</sup> |
| Predator-prey |  |  |  |  |  |  |  |
| Becks et al. (3) | Prey | Cyclic | S - U | U - S | U - NS |  | NS - U |
| Frickel et al. (9) | Both | Stable | S - S | S - S | NS - NS | S - S | NS - S |
| Haafke et al. (13) | Both | Cyclic | U - U | U - U | U - NS | NS - U | NS - U |
| Kasada et al. (26) | Prey | Stable | S - U | U - S | U - NS |  | N - U |
| Wei et al. (34) Fig. 5a | Both | Stable | S - S | S - NS | S - NS | N - S | N - S |
| Fig. 5b | Both | Stable | S - S | S - U | U - NS | N - S | N - U |
| Yoshida et al. (11; 10) | Prey | Cyclic | S - U | U - S | U - NS |  | NS - U |
| Intraguild predation |  |  |  |  |  |  |  |
| Hiltunen et al. (35)<sup>‡</sup> Fig. 2.1b | Basal prey | Cyclic | S - U | U - S | U - S |  | S - U |
| Fig. 2.1c | Basal prey | Cyclic | S - U | U - S | U - U |  | U - U |
| Fig. 2.1d | Basal prey | Cyclic | S - U | U - S | U - S |  | S - U |
| Host-parasite |  |  |  |  |  |  |  |
| Bolker et al. (21) | Parasite | Stable | S - S |  | S - NS | S - S | NS - S |
| Duffy et al. (27) | Host | Stable | S - S | S - S | S - S |  | U - S |

\* The first letter is the stability of the subsystem listed in the column and the second is the stability of the complementary subsystem. S = all eigenvalues for the submatrix have negative real part; U = at least one eigenvalue has positive real part; NS = at least one eigenvalue has negative real part and at least one eigenvalue is zero; N = all eigenvalues are zero. Eco = ecological subsystem; Evo = evolutionary subsystem; Eco-evo = eco-evolutionary subsystem; Pred = exploiter (predator or pathogen); Prey = victim (prey or host).
<sup>†</sup> For systems without prey (predator) evolution, the prey (predator) eco-evolutionary subsystem is just the prey (predator) ecological subsystem.
<sup>‡</sup> Prey subsystems refer to subsystems with just the basal resource variables and predator subsystems refer to subsystems with both the intraguild prey and the intraguild predator variables.

#### S4.1 Becks et al. (3) Model

The discrete trait model is

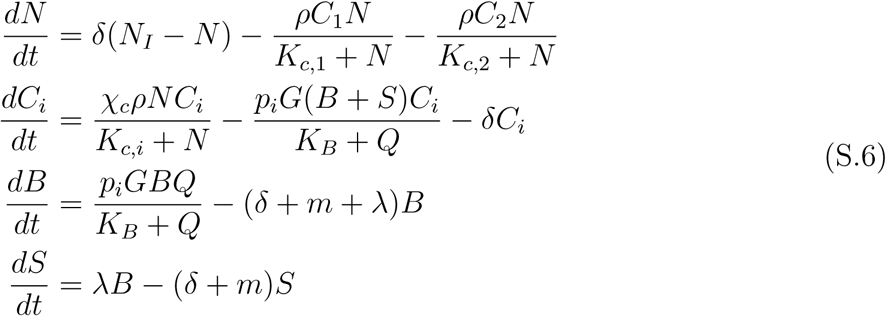

where *N* is the concentration of nitrogen, *C*_*i*_ is the density of algal clones (*i* = 1, 2), *B* and *S* are the densities of breeding and senescent rotifers, and *Q* = *p*_1_*C*_1_ + *p*_2_*C*_2_. The parameter values are *p*_2_ = 1, *N*_*I*_ = 160, *δ* = 0.3, *m* = 0.055*e, λ* = 0.4, *χ*_*c*_ = 0.0027, *K*_*c*,2_ = 2.2, *K*_*c*,1_ = 8 − 5.8*p*_1_, *ρ* = 270, *χ*_*B*_ = 170, *K*_*B*_ = 0.15, *G* = 0.011, *p*_1_ = 0.1.

The four ecological variables for the continuous trait model are *N, C* = *C*_1_+*C*_2_, *B*, and *S*, and the single evolutionary variable is *x*_1_ = *C*_1_*/C*. The equilibrium of the continuous trait model is (*N, C, B, S, x*_1_) = (5.91, 0.338, 2.49, 2.8, 0.777). The Jacobian for the continuous trait model is

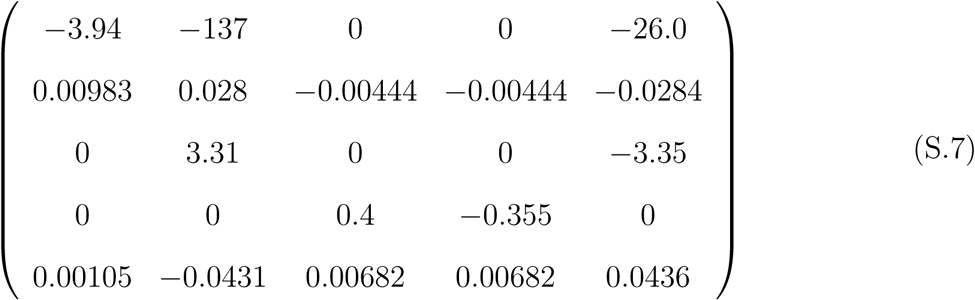

where the order of the rows and columns is (*N, C, B, S, x*_1_). The eigenvalues are (−3.58, −0.462, −0.23, 0.0219±0.229*i*), which implies the equilibrium is unstable. The instability of the equilibrium is due to a Hopf bifurcation that occurs at *δ* ≈ 0.55. The stabilities of the complementary pairs are given in Table S2.

**Table S2:**
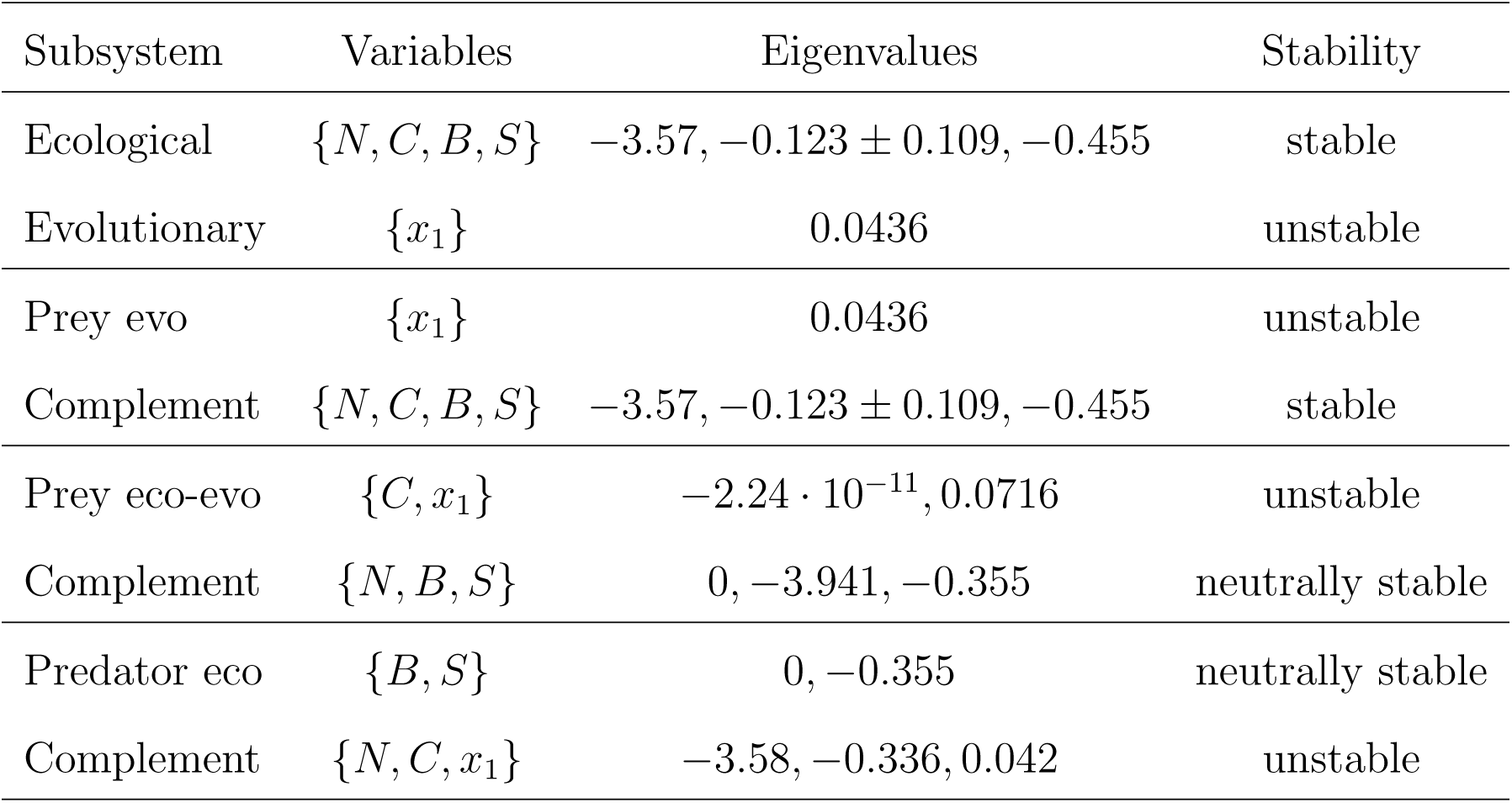
Stabilities of subsystems for the Becks et al. (3) model.

#### S4.2 Bolker et al. (21) Model

The model is

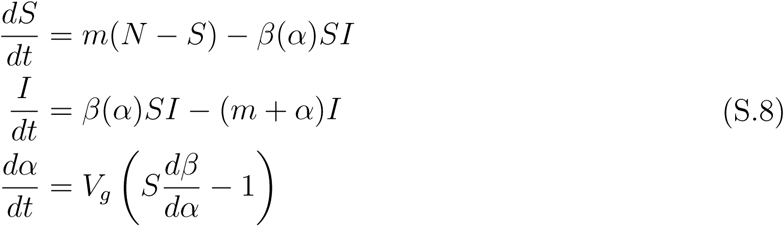

where *S* is the density of susceptible hosts, *I* is the density of infected hosts, and *α* is the population mean virulence of the pathogen; *S* and *I* are ecological variables and *α* is the evolutionary variable. The transmission rate is defined by *β*(*α*) = *cα*^1*/γ*^. Multiple parameterizations are provided for this model because it was applied to four different disease systems. (The values of *c* and *γ* are computed from the reported values of *R*_0_ and *α*^*^; see original study for details). The parameters for SARS were *N* = 1, *m* = 1, *c* = 4.45, *γ* = 1.156. The parameters for HIV were *N* = 1, *m* = 1, *c* = 2.13, *γ* = 1.157. The parameters for West Nile Virus (WNV) were *N* = 1, *m* = 1, *c* = 3.23, *γ* = 1.002. The parameters for Myxomatosis (Myx) were *N* = 1, *m* = 1, *c* = 4.71, *γ* = 1.2.

The equilibrium for the SARS parameterization is (*S, I, α*) = (0.333, 0.00104, 640); the equilibrium for the HIV parameterization is (*S, I, α*) = (.699, 0.0409, 6.36); the equilibrium for the WNV parameterization is (*S, I, α*) = (.309, 0.001082, 639); and the equilibrium for the Myx parameterization is (*S, I, α*) = (1*/*3, 1*/*9, 5). The Jacobians for the different parameterizations are

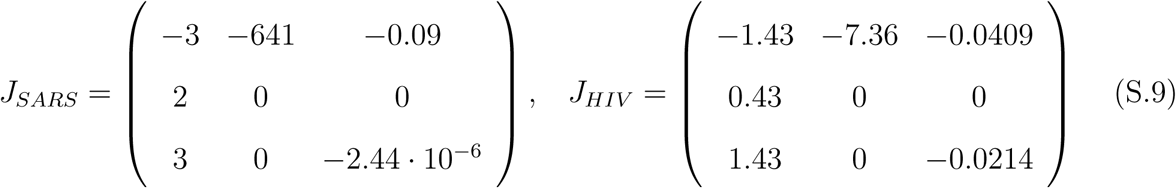

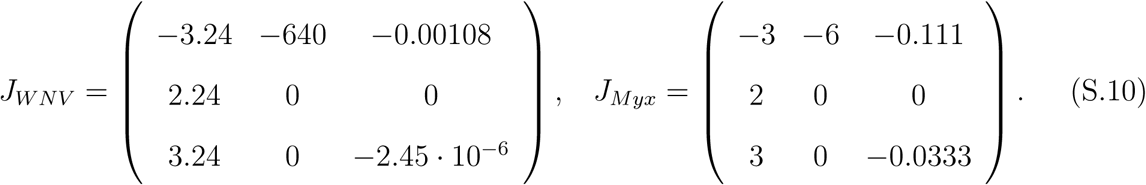

where the orders of the columns and rows are (*S, I, α*). The eigenvalues of *J*_*SARS*_ are (−1.5 ± 35.8*i*, −2.44 · 10^−6^); the eigenvalues of *J*_*HIV*_ are (−0.715 ± 1.65*i*, −0.021); the eigenvalues of *J*_*W NV*_ are (−1.62 ± 37.8*i*, −2.45 · 10^−6^); and the eigenvalues of *J*_*Myx*_ are (−1.5 ± 3.18*i*, −0.0324). In all cases the equilibria are stable. The stabilities of the complementary pairs are given in Table S3. Note that the ecological subsystem {*S, I*} and the complementary evolutionary subsystem is {*α*} also define the parasite evolutionary subsystem and its complement.

**Table S3:** Stabilities of subsystems for the Bolker et al. (21) model.

|  | Subsystem | Variables | Eigenvalues | Stability |
| --- | --- | --- | --- | --- |
| **WNV** | Ecological | $\{S, I\}$ | $-1.5 \pm 35.8i$ | stable |
| | Evolutionary | $\{\alpha\}$ | $-2.44 \cdot 10^{-6}$ | stable |
| | Parasite eco-evo | $\{I, \alpha\}$ | $0, -2.44 \cdot 10^{-6}$ | neutrally stable |
| | Complement | $\{S\}$ | $-3$ | stable |
| **HIV** | Ecological | $\{S, I\}$ | $-0.715 \pm 1.65i$ | stable |
| | Evolutionary | $\{\alpha\}$ | $-0.021$ | stable |
| | Parasite eco-evo | $\{I, \alpha\}$ | $0, -0.021$ | neutrally stable |
| | Complement | $\{S\}$ | $-1.43$ | stable |
| **SARS** | Ecological | $\{S, I\}$ | $-1.62 \pm 37.8i$ | stable |
| | Evolutionary | $\{\alpha\}$ | $-2.45 \cdot 10^{-6}$ | stable |
| | Parasite eco-evo | $\{I, \alpha\}$ | $0, -2.45 \cdot 10^{-6}$ | neutrally stable |
| | Complement | $\{S\}$ | $-3.24$ | stable |
| **Myx** | Ecological | $\{S, I\}$ | $-1.5 \pm 3.18i$ | stable |
| | Evolutionary | $\{\alpha\}$ | $-0.0333$ | stable |
| | Parasite eco-evo | $\{I, \alpha\}$ | $0, -0.0333$ | neutrally stable |
| | Complement | $\{S\}$ | $-3$ | stable |

#### S4.3 Duffy et al. (27) Model

The model is

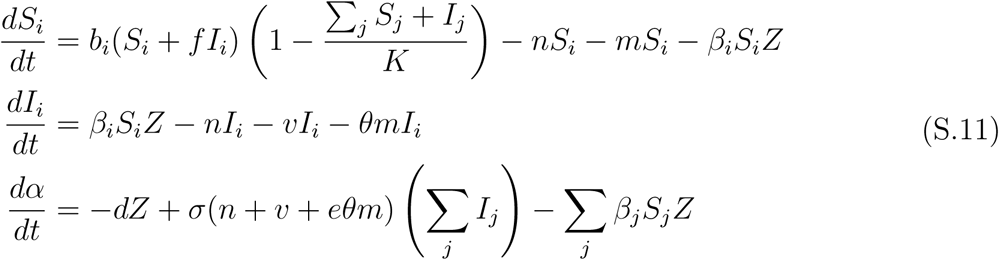

where *S*_*i*_ and *I*_*i*_ are the densities of susceptible and infected clonal hosts (*i* = 1, 2), respectively, and *Z* is the spore density. The parameter values for the model are *b*_1_ = 5712.11*β*_1_ + 0.241, *b*_2_ = 5712.11*β*_2_ + 0.241, *d* = 0.05, *e* = 0.5, *f* = 0.75, *K* = 5, *m* = 0.1, *n* = 0.05, *v* = 0.05, *β*_1_ = 0.5 · 10^−6^, *β*_2_ = 8.5 · 10^−6^, *θ* = 9, *σ* = 15000.

The three ecological variables for the continuous trait model are *S* = *S*_1_ + *S*_2_, *I* = *I*_1_ + *I*_2_, and *Z* and the evolutionary variables are the proportions of host clone one in each stage, *x*_1_ = *S*_1_*/S* and *x*_2_ = *I*_2_*/I*. The equilibrium of the continuous trait model is (*S, I, Z, x*_1_, *x*_2_) = (1.86, 0.025, 4120, 0.656, 0.101). The Jacobian for the continuous trait system is

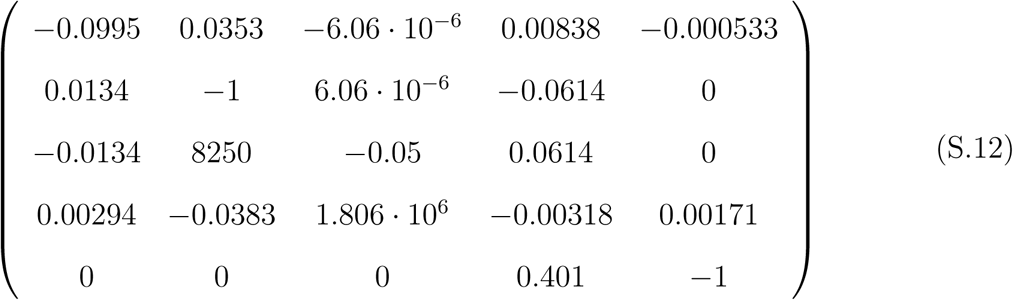

where the order of the rows and columns is (*S, I, Z, x*_1_, *x*_2_). The eigenvalues are (−1.05, −1, −0.0915, −0.00318 ± 0.027*i*), which implies the equilibrium is stable. The stabilities of the complementary pairs are given in Table S4.

**Table S4:**
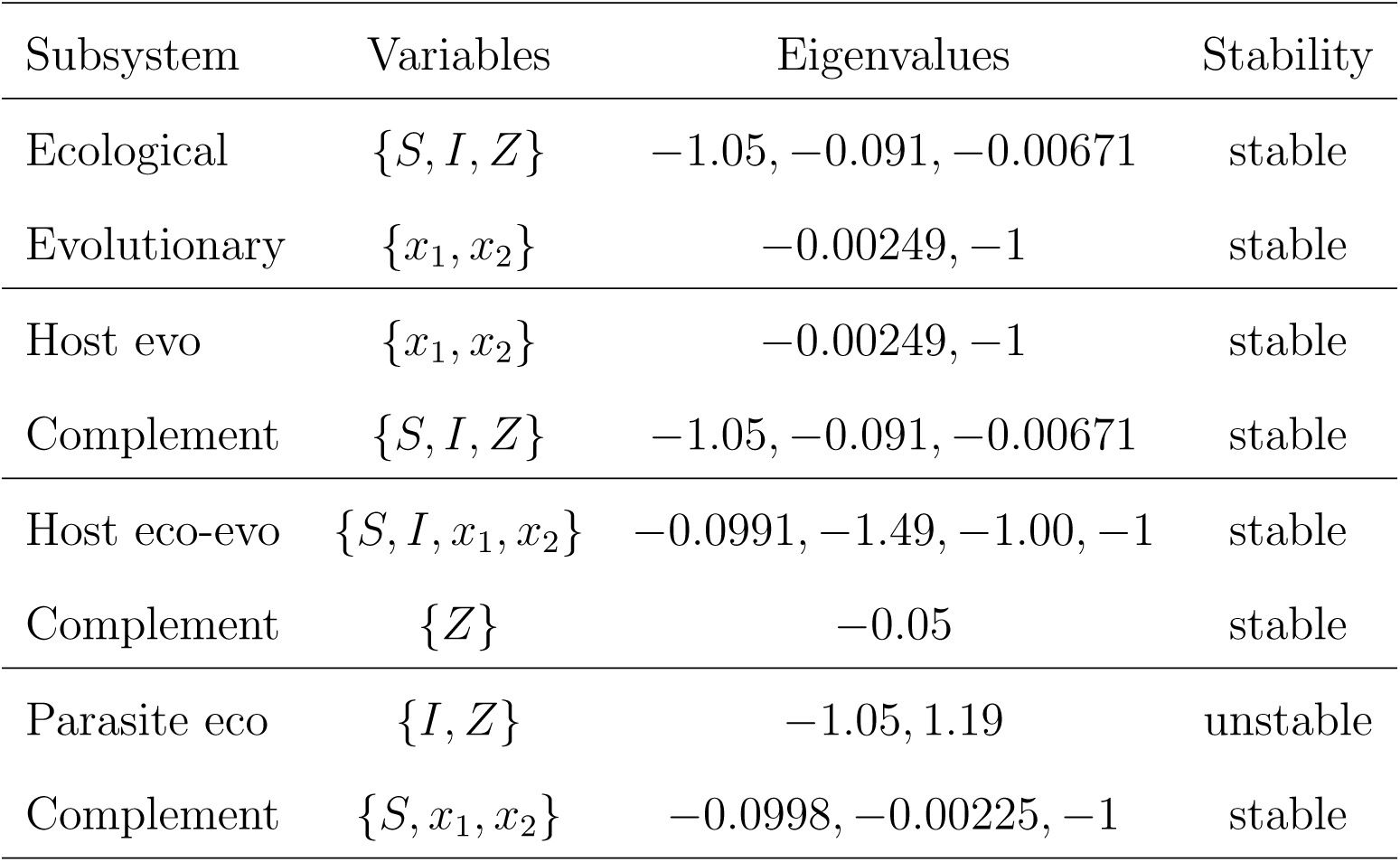
Stabilities of subsystems for the Duffy et al. (27) model.

#### S4.4 Frickel et al. (9) Model

In matrix form the model is

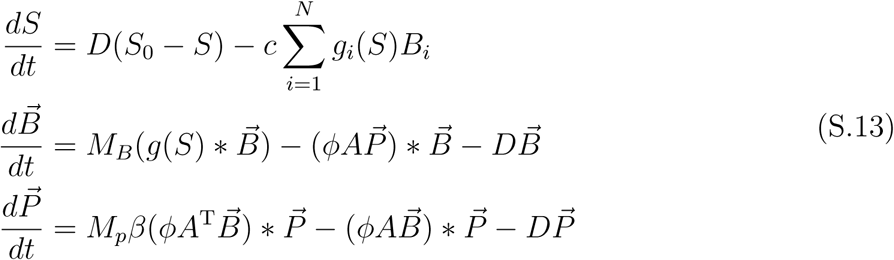

where *S* is the resource concentration, 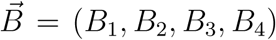 is the density of algal clones, and 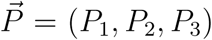 is the density of viral types. In the model, * denotes component-wise multiplication, *g*_*i*_(*S*) = *a*_*i*_*S/*(*H* + *S*), *a*_*i*_ = *a*_1_ + (*a*_*N*_ − *a*_1_)(*i* − 1)*/*(*N* − 1), and *A* is an upper triangular 4×3 matrix with ones on and above the main diagonal. The matrices *M*_*B*_ and *M*_*p*_are

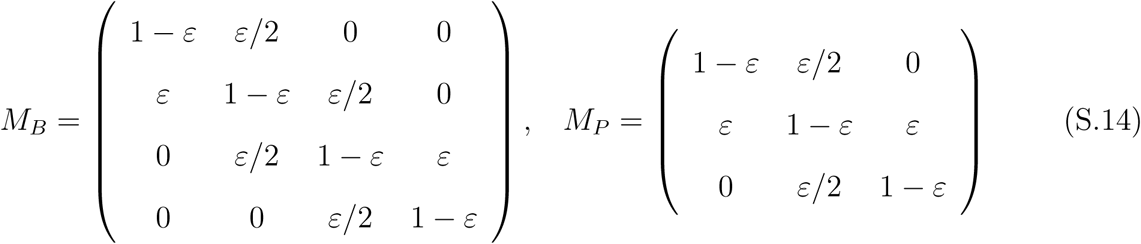

The parameter values are *a*_1_ = 0.25, *a*_*N*_ = 0.15, *D* = 0.1, *S*_0_ = 30, *H* = 1, *c* = 2.3 · 10^−5^, *ϕ* = 7.5 · 10^−8^, *β* = 100, *ε* = 10^−3^.

The three ecological variables for the continuous trait model are *S*, total algal density (*B* = *B*_1_ + *B*_2_ + *B*_3_ + *B*_4_), and total viral density (*P* = *P*_1_ + *P*_2_ + *P*_3_). The evolutionary variables are *x*_1_ = *B*_1_*/B, x*_2_ = *B*_2_*/B, x*_3_ = *B*_3_*/B, x*_4_ = *P*_1_*/P, x*_5_ = *P*_2_*/P*. The equilibrium of the continuous trait model is (*S, B, P, x*_1_, *x*_2_, *x*_3_, *x*_4_, *x*_5_) = (2.01, 1210000, 0.00884, 0.726 · 10^−4^, 0.00224, 889000, 0.973 · 10^−5^, 0.005). The Jacobian is

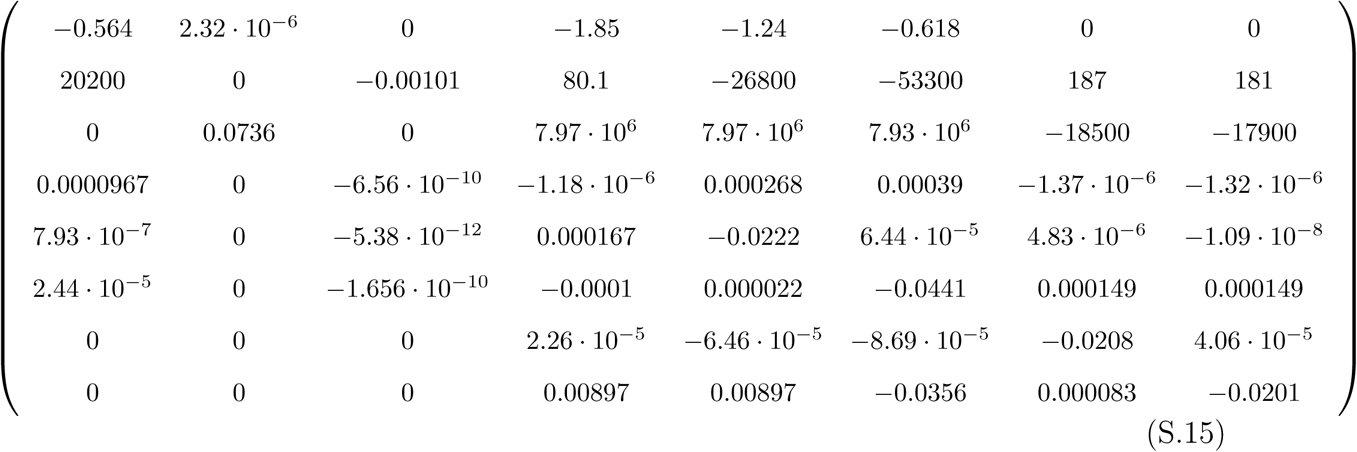

where the order of the columns and rows is (*S, B, P, x*_1_, *x*_2_, *x*_3_, *x*_4_, *x*_5_). The eigenvalues are (−0.463, −0.00437 ± 0.0792*i*, −0.1, −0.0365, −0.0204, −0.0207, −0.0222), which implies the equilibrium is stable. The stabilities of the complementary pairs are given in Table S5.

**Table S5:**
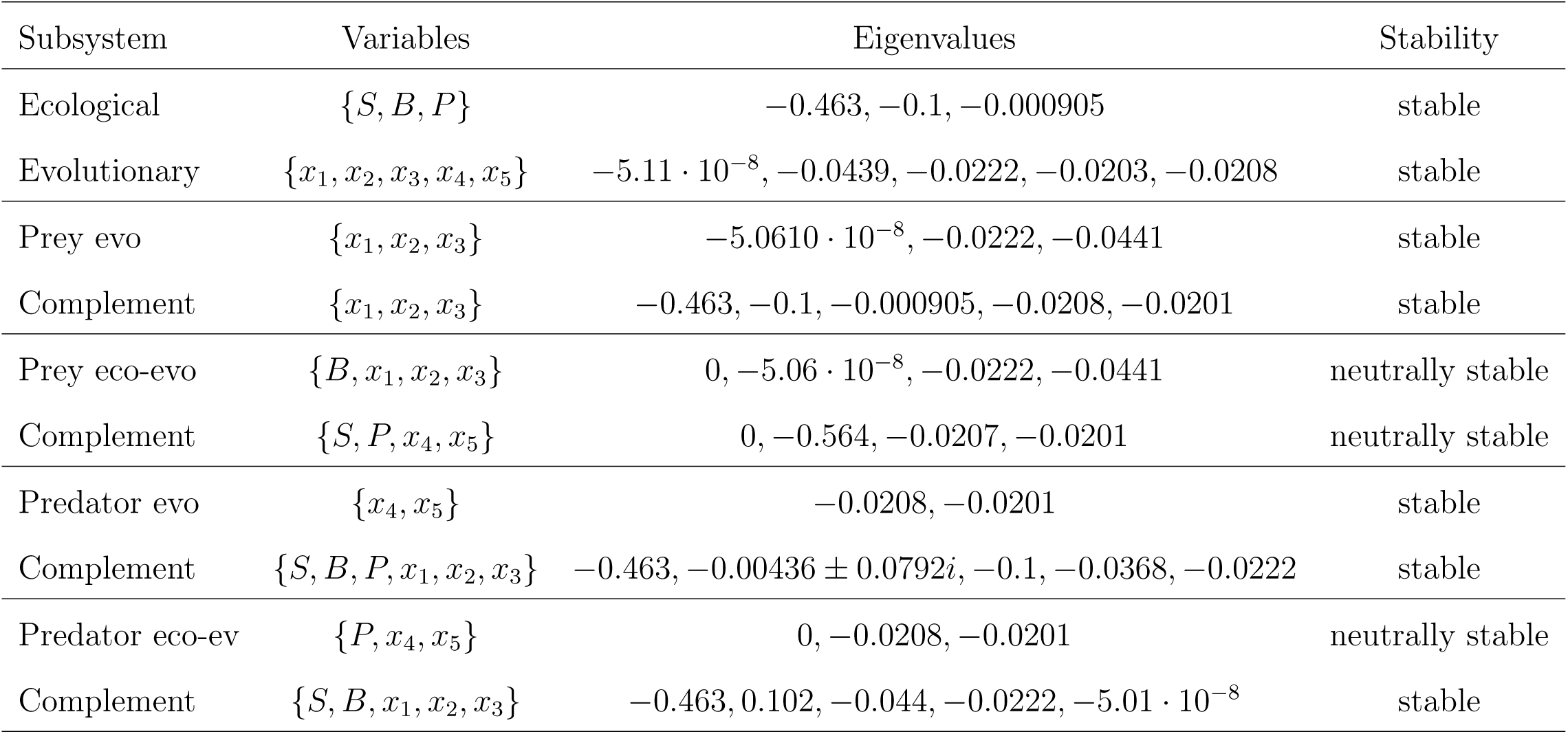
Stabilities of subsystems for the Frickel et al. (9) model.

#### S4.5 Haafke et al. (13) Model

The model is

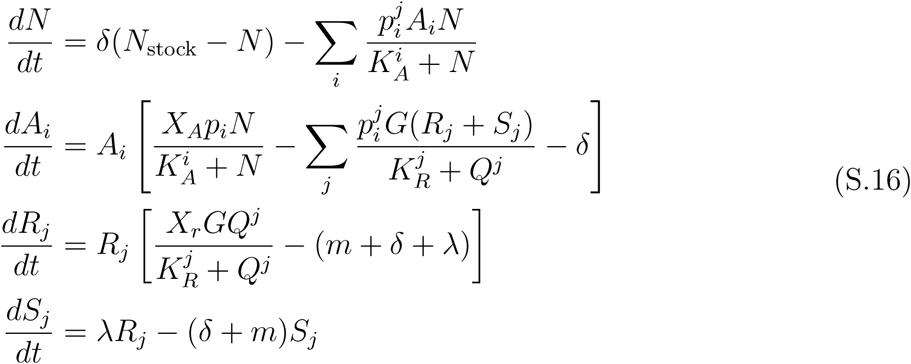

where *N* is the concentration of nitrogen, *A*_*i*_ is the density of algal clones (*i* = 1, 2), *R*_*j*_ and *S*_*j*_ are the densities of breeding and senescent rotifer clones (*j* = 1, 2), and 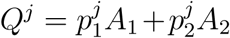. The parameter values are 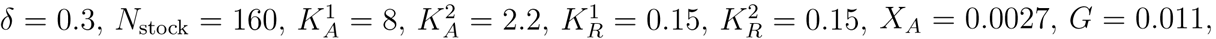 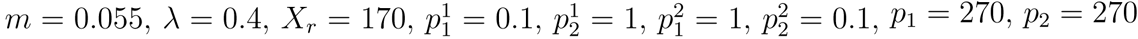.

The four ecological variables of the continuous trait model are nitrogen concentration (*N*), total algal density (*A* = *A*_1_ + *A*_2_), and total breeding (*R* = *R*_1_ + *R*_2_) and senescent (*S* = *S*_1_ + *S*_2_) rotifers. The evolutionary variables are *x*_1_ = *A*_1_*/A, x*_2_ = *R*_1_*/R*, and *x*_3_ = *S*_1_*/S*. The equilibrium of the continuous trait model is (*N, A, R, S, x*_1_, *x*_2_, *x*_3_) = (22.7, 0.185, 5.9, 6.65, 0.5, 0.627, 0.627). The Jacobian is

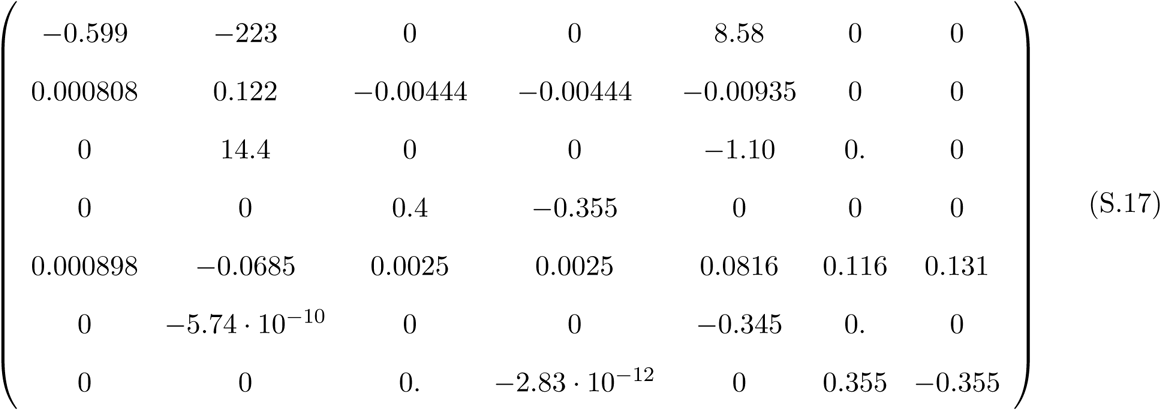

where the order of the columns and rows is (*N, A, R, S, x*_1_, *x*_2_, *x*_3_). The eigenvalues are (0.0156 ± 0.369*i*, 0.0602 ± 0.274*i*, −0.417 ± 0.133*i*, −0.423), which implies the equilibrium is unstable. The instability of the equilibrium is due to a Hopf bifurcation that occurs at *δ* ≈ 0.734. The stabilities of the complementary pairs are given in Table S6.

**Table S6:**
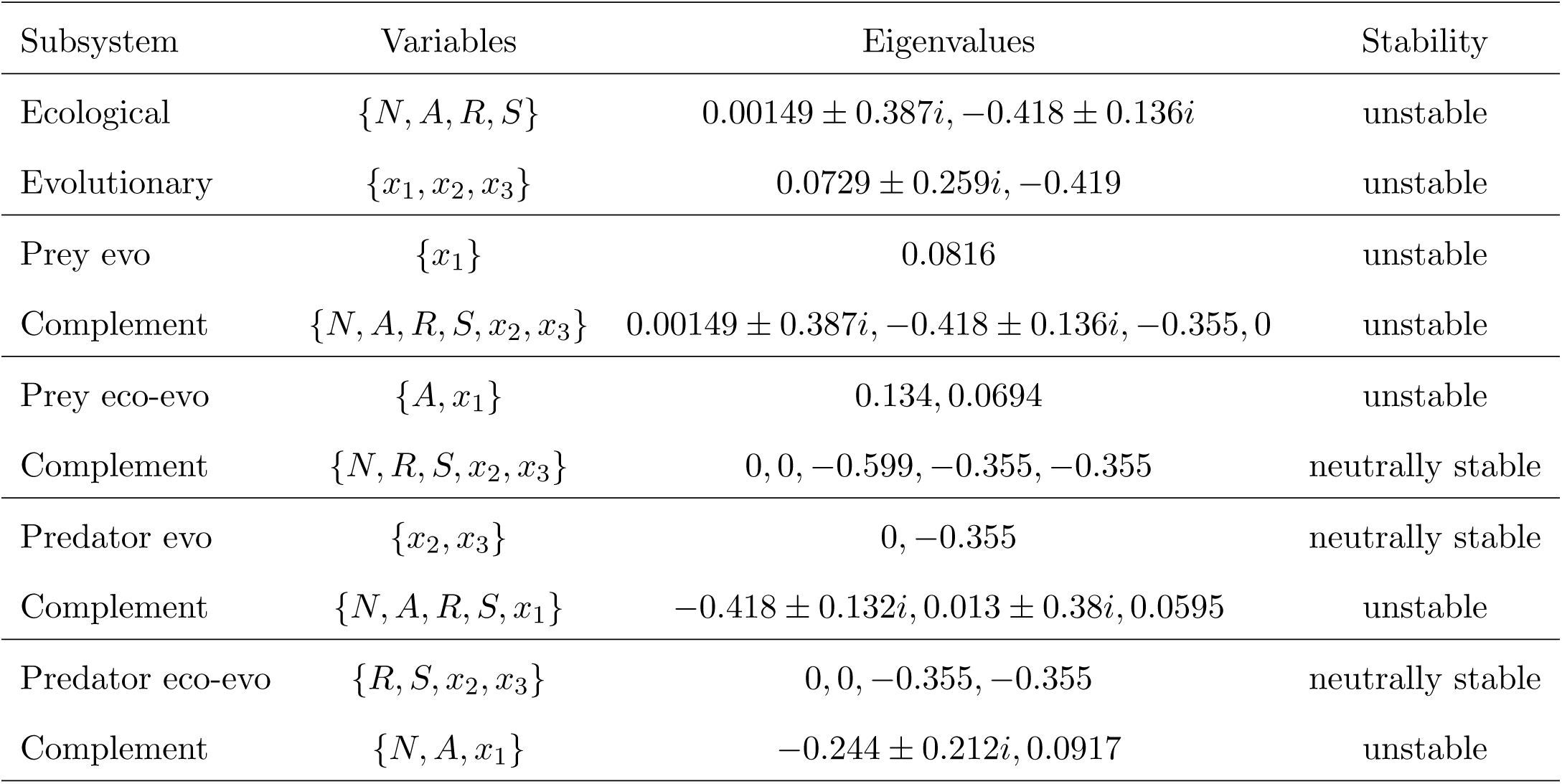
Stabilities of subsystems for the Haafke et al. (13) model.

#### S4.6 Hiltunen et al. (35) Model

The model is

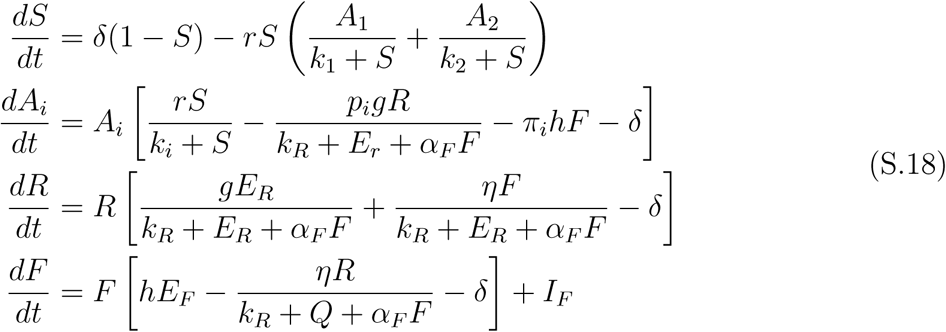

where *S* is the concentration of limiting substrate, *A*_*i*_ is the density of algal clones (*i* = 1, 2), *R* is the density of rotifers (the intraguild predator), and *F* is the density of flagellates (the intraguild prey). In the model *E*_*R*_ = *p*_1_*A*_1_ + *p*_2_*A*_2_ and *E*_*F*_ = *π*_1_*A*_1_ + *π*_2_*A*_2_. The three parameterizations for figures 1b-d are listed below.

The four ecological variables of the continuous trait model are *S, A* = *A*_1_ + *A*_2_, *R*, and *F* and the evolutionary variable is *x*_1_ = *A*_1_*/A*. Note that the ecological subsystem {*S, A, R, F*} and the complementary evolutionary subsystem {*x*_1_} also define the prey evolutionary subsystem and its complement.

**Figure 1b**: The parameter values are *δ* = 1, *r* = 2, *k*_1_ = 0.234, *k*_2_ = 0.19, *p*_1_ = 0.05, *p*_2_ = 1, *g* = 2, *k*_*r*_ = 0.2, *α*_*F*_ = 0.05, *π*_1_ = 1, *π*_2_ = 0.1, *h* = 3, *η* = 0.08, *I*_*F*_ = 0.001. The equilibrium of the continuous trait model is (*S, A, R, F, x*_1_) = (0.372, 0.497, 0.0608, 0.0711, 0.633). The Jacobian for the continuous trait model is

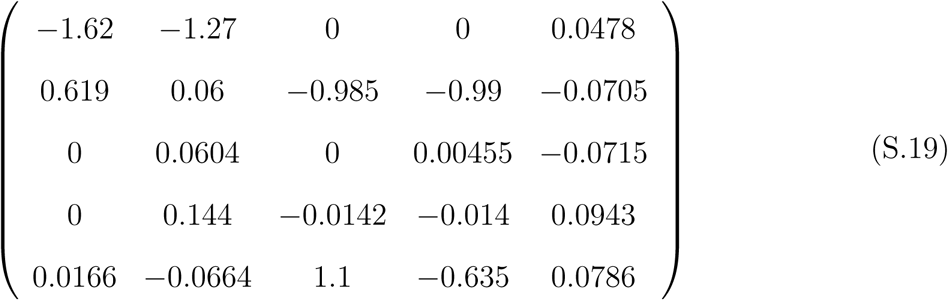

where the order of the columns and rows is (*S, A, R, F, x*). The eigenvalues are (−1, −0.29 ± 0.506*i*, 0.0422 ± 0.363*i*), which implies the equilibrium is unstable. The instability of the equilibrium is due to a Hopf bifurcation that occurs at *δ* ≈ 1.17. The stabilities of the complementary pairs are given in Table S7.

**Figure 1c**: The parameter values are *δ* = 1, *r* = 3.3, *k*_1_ = 0.1475, *k*_2_ = 0.07375, *p*_1_ = 0.05, *p*_2_ = 1, *g* = 2.5, *k*_*R*_ = 0.2, *α*_*F*_ = 0.05, *π*_1_ = 1, *π*_2_ = 0.05, *h* = 3, *η* = 0.4, *I*_*F*_ = 0.001. The equilibrium of the continuous trait model is (*S, A, R, F, x*_1_) = (0.168, 0.456, 0.146, 0.231, 0.869). The Jacobian for the continuous trait model is

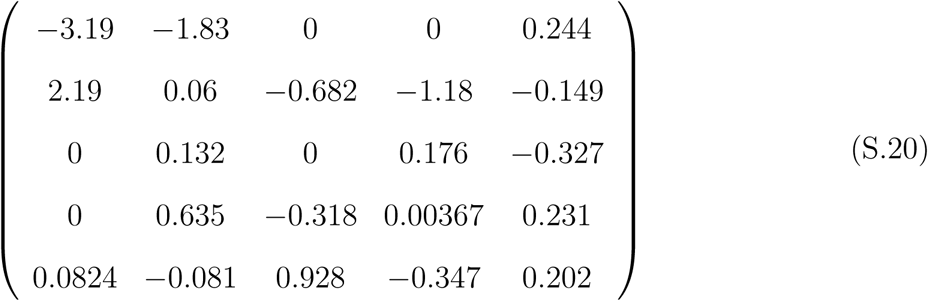

where the order of the columns and rows is (*S, A, R, F, x*). The eigenvalues are (−1 ± 1.12, −1, 0.0395 ± 0.619*i*), which implies the equilibrium is unstable. The instability of the equilibrium is due to a Hopf bifurcation that occurs at *δ* ≈ 1.69. The stabilities of the complementary pairs are given in Table S7.

**Figure 1d**: The parameter values are *δ* = 1, *r* = 2, *k*_1_ = 0.19, *k*_2_ = 0.12, *p*_1_ = 0.1, *p*_2_ = 1, *g* = 2, *k*_*R*_ = 0.2, *α*_*F*_ = 0.05, *π*_1_ = 1, *π*_2_ = 1, *h* = 3, *η* = 0.4, *I*_*F*_ = 0.001. The equilibrium of the continuous trait model is (*S, A, R, F, x*_1_) = (0.485, 0.343, 0.03307748540, 0.14, 0.622). The Jacobian for the continuous trait model is

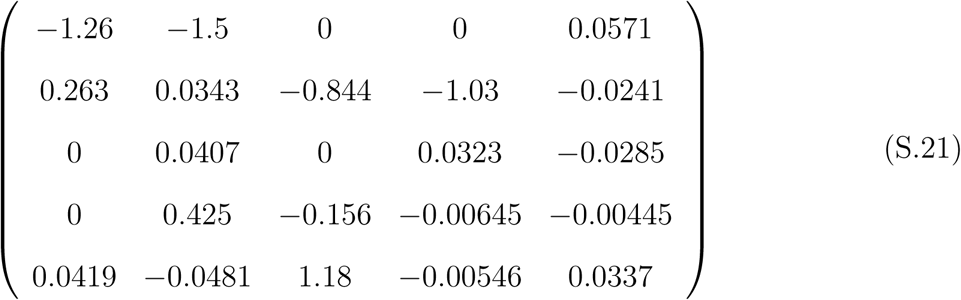

where the order of the columns and rows is (*S, A, R, F, x*). The eigenvalues are(−1, −0.106 ± 0.763*i*, 0.0052 ± 0.174*i*), which implies the equilibrium is unstable. The instability of the equilibrium is due to a Hopf bifurcation that occurs at *δ* ≈ 1.17. The stabilities of the complementary pairs are given in Table S7.

**Table S7:** Stabilities of subsystems for the Hiltunen et al. (35) model.

|  | Subsystem | Variables | Eigenvalues | Stability |
| --- | --- | --- | --- | --- |
| **Fig. 1b** | Ecological | $\{N, C, B, S\}$ | $-1, -0.285 \pm 0.503i, -0.00317$ | stable |
| | Evolutionary | $\{x_1\}$ | 0.0786 | unstable |
| | Prey eco-evo | $\{C, x_1\}$ | $-6.32 \cdot 10^{-11}, 0.138$ | unstable |
| | Complement | $\{N, B, S\}$ | $-1.62, -0.00698 \pm 0.00395i$ | stable |
| | Predator eco | $\{B, S\}$ | $-0.00698 \pm 0.00395i$ | stable |
| | Complement | $\{N, C, x_1\}$ | $-0.784 \pm 0.283i, 0.0877$ | unstable |
| **Fig. 1c** | Ecological | $\{N, C, B, S\}$ | $-0.997 \pm 1.146i, -1, -0.129$ | stable |
| | Evolutionary | $\{x_1\}$ | 0.202 | unstable |
| | Prey eco-evo | $\{C, x_1\}$ | $-1.93 \cdot 10^{-11}, 0.262$ | unstable |
| | Complement | $\{N, B, S\}$ | $-3.19, 0.00183 \pm 0.237i$ | unstable |
| | Predator eco | $\{B, S\}$ | $0.00183 \pm 0.237i$ | unstable |
| | Complement | $\{N, C, x_1\}$ | $-1.57 \pm 1.16i, 0.207$ | unstable |
| **Fig. 1d** | Ecological | $\{N, C, B, S\}$ | $-1, -0.11 \pm 0.764i, -0.0142$ | stable |
| | Evolutionary | $\{x_1\}$ | 0.0337 | unstable |
| | Prey eco-evo | $\{C, x_1\}$ | $0.068, 1.34 \cdot 10^{-12}$ | unstable |
| | Complement | $\{N, B, S\}$ | $-1.263, -0.00322 \pm 0.0709i$ | stable |
| | Predator eco | $\{B, S\}$ | $-0.00322 \pm 0.0709i$ | stable |
| | Complement | $\{N, C, x_1\}$ | $-0.779, -0.46, 0.0395$ | unstable |

#### S4.7 Kasada et al. (26) Model

The model is

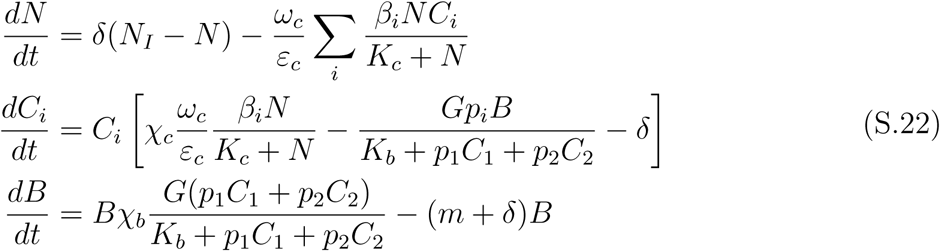

where *N* is concentration of nitrogen, *C*_*i*_ is the density of algal clones (*i* = 1, 2), and *B* is the density of rotifers. We only focus on the parameter values for the UTEX396 and UTEX265 algal clones because coexistence of multiple clonal types did not occur for other combinations. This corresponds to figure 4E in Kasada et al. (26). The parameter values are *β*_1_ = 1.77, *β*_2_ = 1.57, *p*_1_ = 0.102, *p*_2_ = 0.102 · 0.688, *N*_*I*_ = 80, *δ* = 0.5, *χ*_*c*_ = 0.05, *χ*_*b*_ = 54000, *m* = 0.055, *K*_*c*_ = 4.3, *K*_*b*_ = 0.835, *ω*_*c*_ = 20, *ε* = 1, *G* = 5 · 10^−5^.

For the continuous trait model the ecological variables are *N, C* = *C*_1_ + *C*_2_, and *B* and the evolutionary variable is *x*_1_ = *C*_1_*/C*. The equilibrium of the continuous trait model is (*N, C, B, x*_1_) = (3.42, 2.63, 58500, 0.38). The Jacobian for the system is

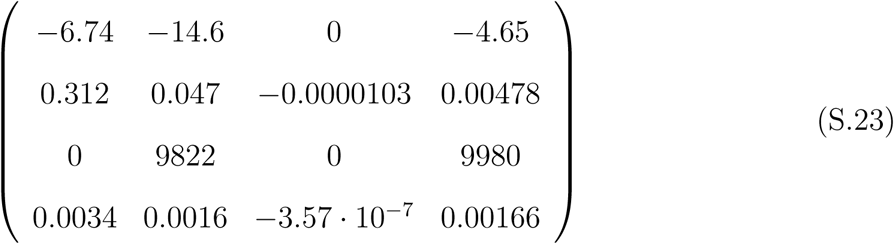

where the order of the columns and rows is (*N, C, B, x*). The eigenvalues are (−5.99, −0.455, −0.239, −0.0117), which implies the equilibrium is stable. The stabilities of the complementary pairs are given in Table S8. Note that the ecological subsystem and the complementary evolutionary subsystem also define the prey evolutionary subsystem and its complement.

**Table S8:**
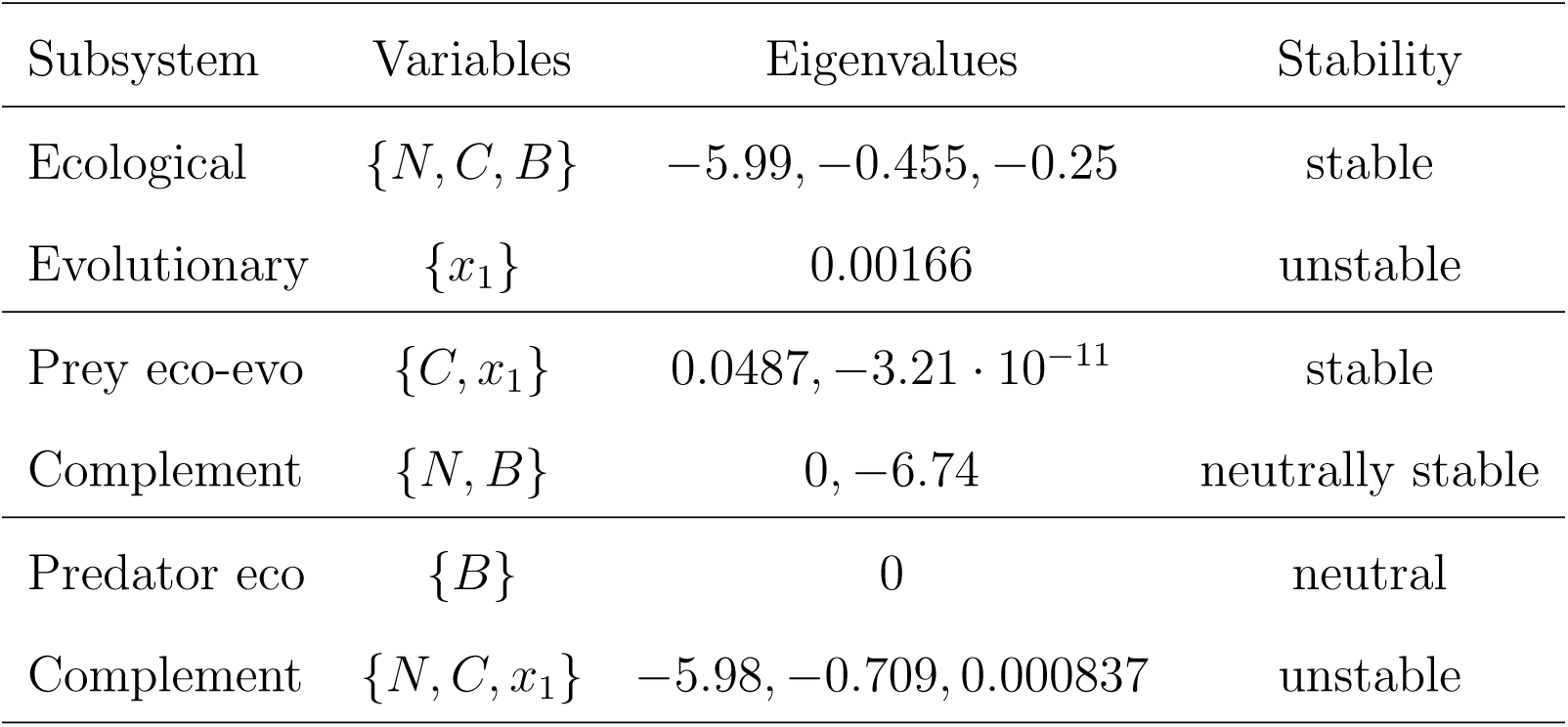
Stabilities of subsystems for the Kasada et al. (26) model.

#### S4.8 Wei et al. (34) Model

The model is

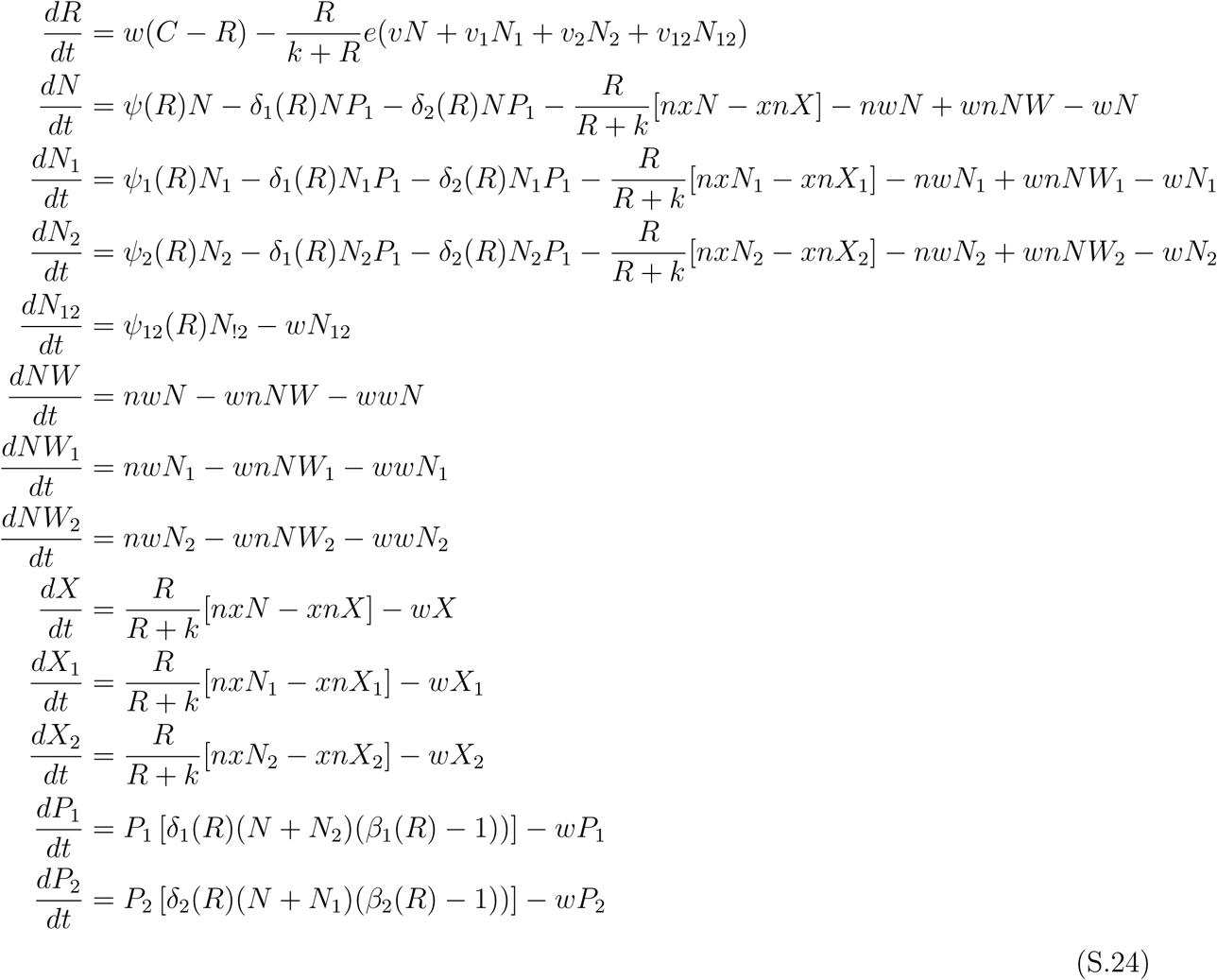

where the variables are phage densities (*P*_1_, *P*_2_), planktonic, non-persistor bacteria densities (*N, N*_1_, *N*_2_, *N*_12_), persistor bacteria densities (*X*_1_, *X*_2_, *X*_3_) and wall bacteria densities (*NW, NW*_1_, *NW*_2_); the subscripts for the bacteria populations denote which phage the bacteria is resistant to. The functions in the model are

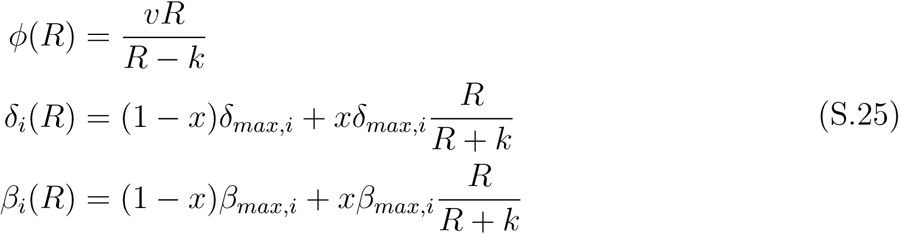

The parameter values are *v* = 1, *k* = 0.25, *e* = 5 · 10^−7^, *δ*_*max*,1_ = 0.1 · 10^−7^, *δ*_*max*,2_ = 0.1 · 10^8^, *β*_*max*,1_ = 50, *β*_*max*,2_ = 100, *x* = 0.5, *w* = 0.4, *ww* = 0.01, *C* = 100, *nx* = 0.0001, *xn* = 0.0001, *nw* = 0.01, *wn* = 0.005, *v*_1_ = 0.9, *v*_2_ = 0.85, *v*_12_ = 0.8.

**Figure 5a:** This parameterization assumes *N, X, NW*, and *N*_12_ are not present, i.e., there are no bacteria susceptible to both or none of the phage types. The ecological variables of the continuous trait model are the total densities of the different bacterial types and phage: 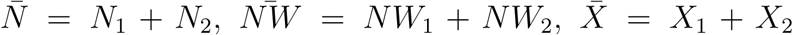, and 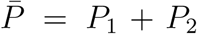. To avoid notational confusion, we use *χ*_*i*_ to denote the evolutionary traits. The evolutionary variables are 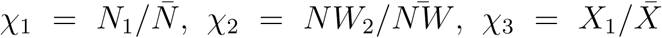, and 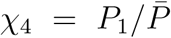. The equilibrium is 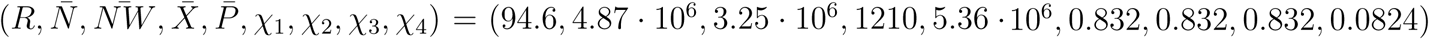. The Jacobian for the system is

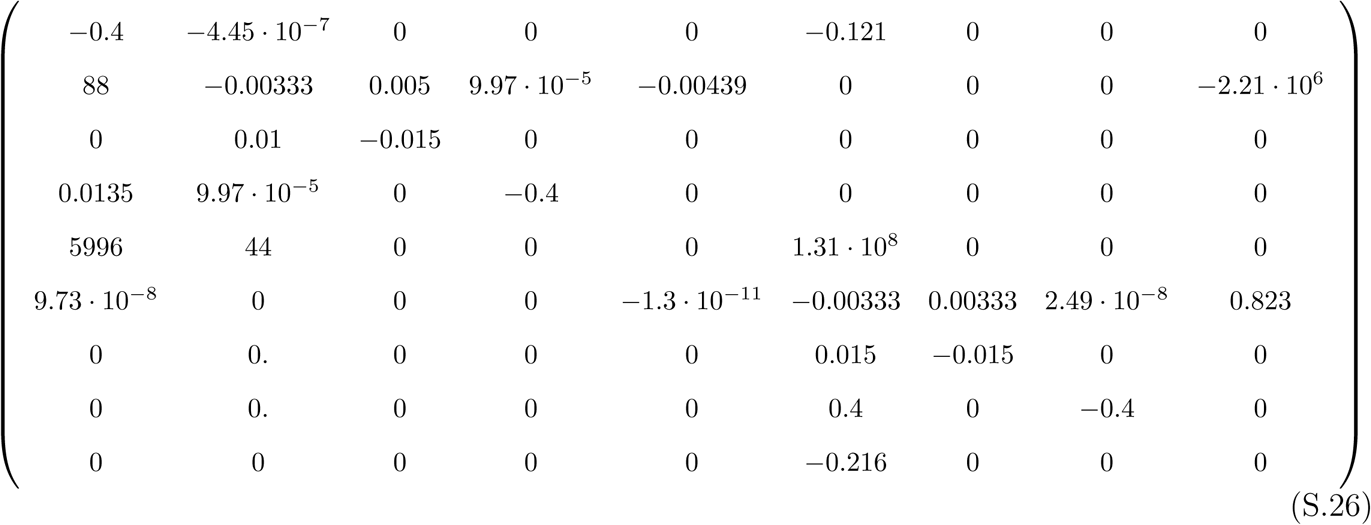

where the order of the columns and rows is 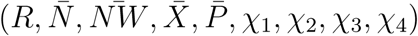. The eigenvalues are (−0.0017 ± 0.443*i*, −0.00167 ± 0.42*i*, −0.015, −0.015, −0.4, −0.4, −0.4), which implies the equilibrium is stable. The stabilities of the complementary pairs are given in Table S9.

**Table S9:**
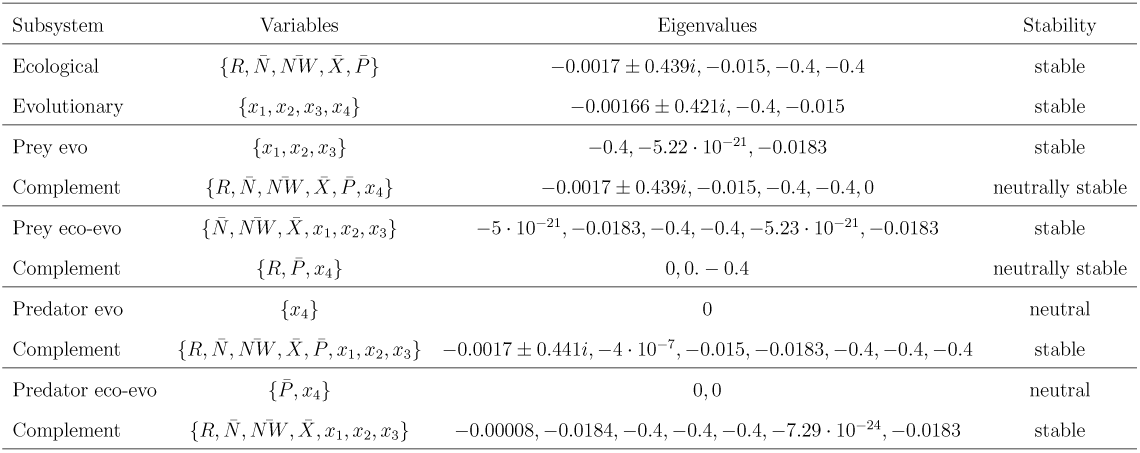
Stabilities of subsystems for the Fig. 5a Wei et al. (34) model.

**Figure 5b:** This parameterization assumes *N*_2_, *X*_2_ and *NW*_2_ are not present, i.e., there are no types that are just resistant to phage 2. For the continuous trait model the ecological variables are the total densities of the different bacterial types and phage: 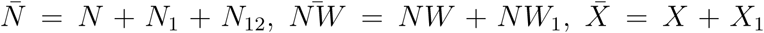, and 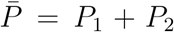. To avoid notational confusion, we use *χ*_*i*_ to denote the evolutionary traits. The evolutionary variables are 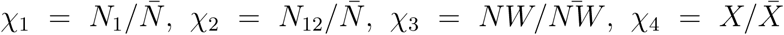 and 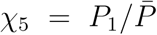. The equilibrium is 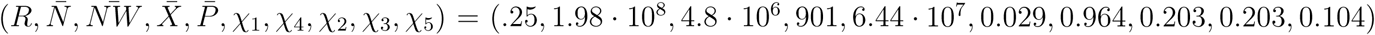. The Jacobian for the system is

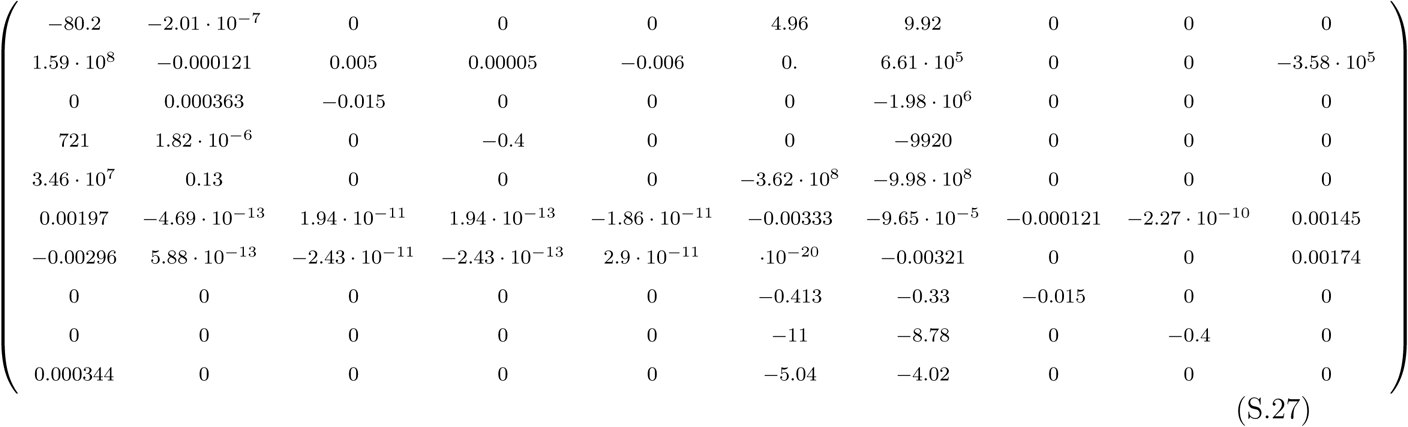

where the order of the columns and rows is 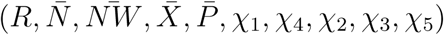. The eigen-values are (−79.8, −0.00198 ± 0.162*i*, −0.00168 ± 0.1*i*, −0.015, −0.0151, −0.401, −0.4 − 0.4), which implies the equilibrium is stable. The stabilities of the complementary pairs are given in Table S10.

**Table S10:** Stabilities of subsystems for the Fig. 5b Wei et al. (34) model.

| Subsystem | Variables | Eigenvalues | Stability |
| --- | --- | --- | --- |
| Ecological | $\{R, \bar{N}, N\bar{W}, \bar{X}, \bar{P}\}$ | $-79.8, -0.000645, -0.015, -0.401, -0.4$ | stable |
| Evolutionary | $\{x_1, x_2, x_3, x_4, x_5\}$ | $-0.4, -0.00164 \pm 0.119i, -0.00321, -0.015$ | stable |
| Prey evo | $\{x_1, x_2, x_3, x_4\}$ | $-0.4, -0.0183, -1.88 \cdot 10^{-18}, -0.00321$ | stable |
| Complement | $\{R, \bar{N}, N\bar{W}, \bar{X}, \bar{P}, x_5\}$ | $-79.8, -0.401, -0.4, -0.015, -0.000645, 3.57 \cdot 10^{-15}$ | unstable |
| Prey eco-evo | $\{\bar{N}, N\bar{W}, \bar{X}, x_1, x_2, x_3, x_4\}$ | $1.6 \cdot 10^{-20}, -0.0183, -0.4, -0.0183$<br>$-0.4, 8.67 \cdot 10^{-20}, 9.31 \cdot 10^{-20}$ | unstable |
| Complement | $\{R, \bar{P}, x_5\}$ | $0, 0, -80.2$ | neutrally stable |
| Predator evo | $\{x_5\}$ | $0$ | neutral |
| Complement | $\{R, \bar{N}, N\bar{W}, \bar{X}, \bar{P}, x_1, x_2, x_3, x_4\}$ | $-79.8, -0.00202 \pm 0.148i, -0.401, -0.4$<br>$-0.4, -0.0183, -0.015, 3.54 \cdot 10^{-16}$ | stable |
| Predator eco-evo | $\{\bar{P}, x_5\}$ | $0, 0$ | neutral |
| Complement | $\{R, \bar{N}, N\bar{W}, \bar{X}, x_1, x_2, x_3, x_4\}$ | $-79.8, -0.0182, -0.402, -0.4$<br>$-0.4, -0.0183, -8.42 \cdot 10^{-20}, 3.02 \cdot 10^{-18}$ | unstable |

#### S4.9 Yoshida et al. (10; 11) Model

The model is

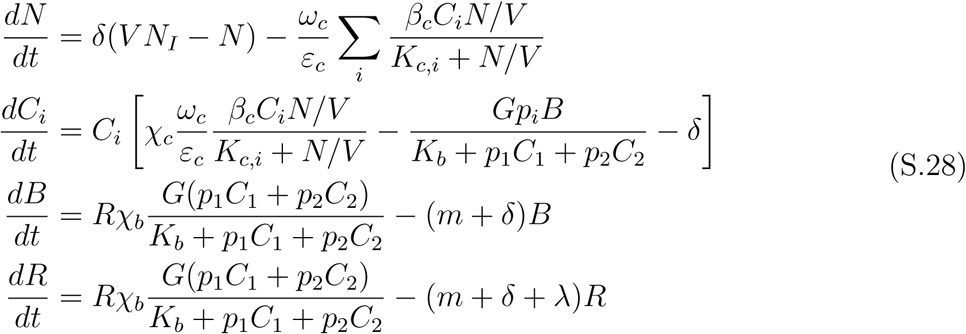

where *N* is concentration of nitrogen, *C*_*i*_ is the density of algal clones (*i* = 1, 2), and *B* and *R* are the total rotifer density and density of fertile rotifers, respectively. In the model 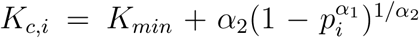. The parameters are *N*_*I*_ = 80, *V* = 0.33, *K*_*min*_ = 4.3, *K*_*b*_ = 0.292, *β*_*c*_ = 3.3, *α*_1_ = 0.8, *α*_2_ = 9.5, *p*_*min*_ = 0.02, *δ* = 0.69, *χ*_*b*_ = 5400, *χ*_*c*_ = 0.05, *ω*_*c*_ = 20, *ε*_*c*_ = 1, *G* = 3.3 · 10^−4^, *λ* = 0.4, *m* = 0.055, *p*_1_ = 0.02, *p*_2_ = 1.

For the continuous trait model the ecological variables are *N, C* = *C*_1_ + *C*_2_, *B*, and *R* and the evolutionary variable is *x*_1_ = *C*_1_*/C*. The equilibrium of the continuous trait model is (*N, C, B, R, x*_1_) = (0.731, 1.07, 690, 449, 0.495). The Jacobian for the system is

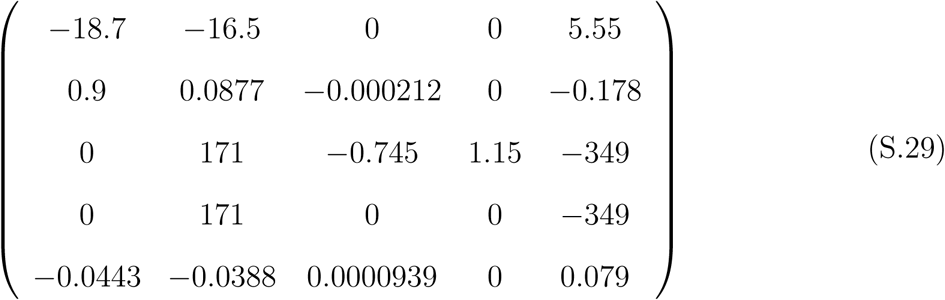

where the order of the columns and rows is (*N, C, B, R, x*_1_). The eigenvalues are (−17.8, −0.864, −0.562, 0.0028 ± 0.219*i*), which implies the equilibrium is unstable. The instability of the equilibrium is due to a Hopf bifurcation that occurs at *δ* ≈ 0.7. The stabilities of the complementary pairs are given in Table S11.

**Table S11:**
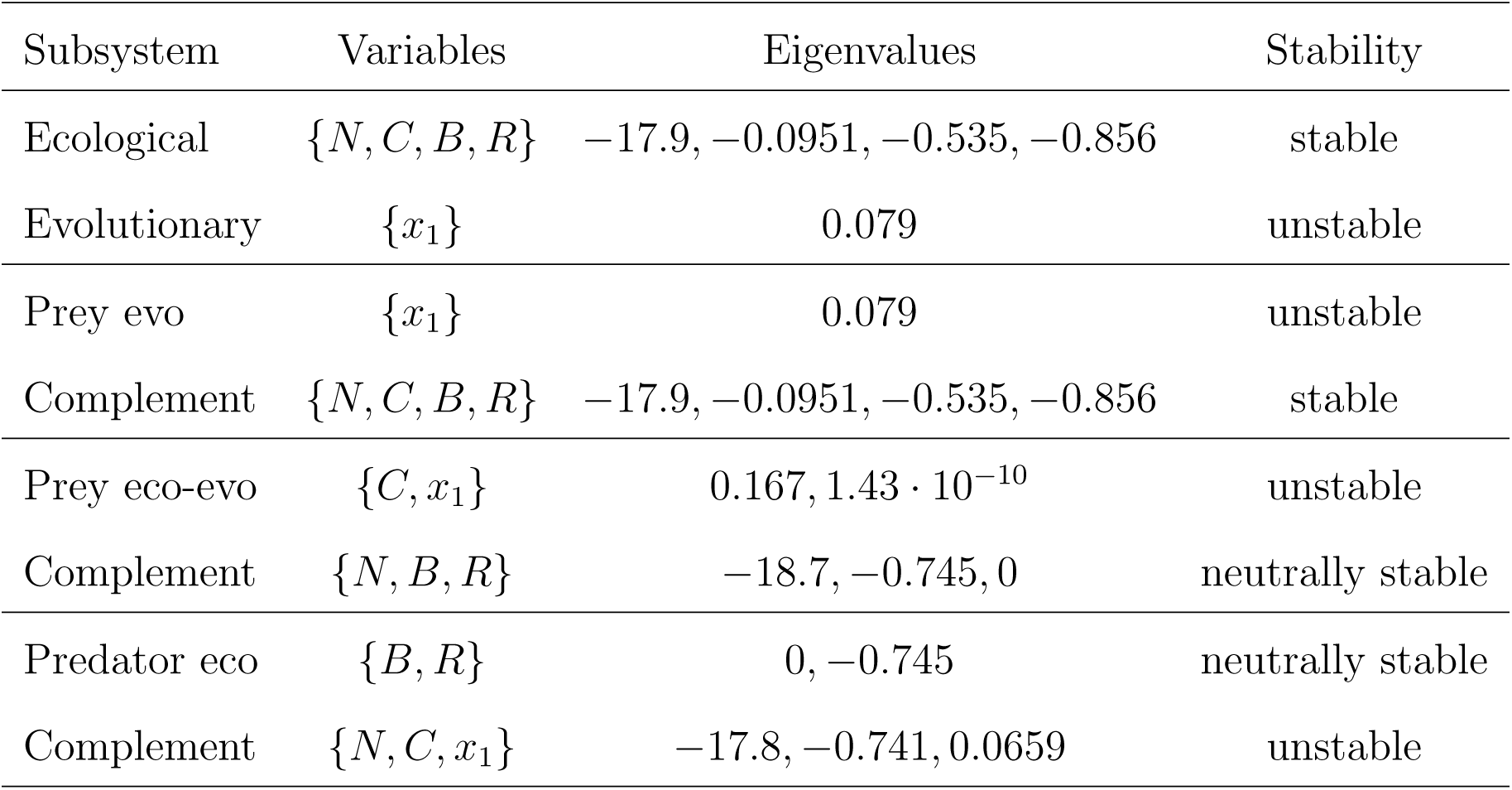
Stabilities of subsystems for the Yoshida et al. (10; 11) model.

